# Cleavage of the TrkB-FL Receptor During Epileptogenesis: Insights from a Kainic Acid-Induced Model of Epilepsy and Human Samples

**DOI:** 10.1101/2024.12.18.628660

**Authors:** Leonor Ribeiro-Rodrigues, João Fonseca-Gomes, Sara L. Paulo, Ricardo Viais, Filipa F. Ribeiro, Catarina Miranda-Lourenço, Francisco M. Mouro, Rita F. Belo, Catarina B. Ferreira, Sara R. Tanqueiro, Mafalda Ferreira-Manso, Juzoh Umemori, Eero Castrén, Vítor H. Paiva, Ana M. Sebastião, Eleonora Aronica, Alexandre Rainha Campos, Carla Bentes, Sara Xapelli, Maria José Diógenes

## Abstract

Brain-derived neurotrophic factor (BDNF) is essential for neuronal survival, differentiation, and plasticity. In epilepsy, BDNF exhibits a dual role, exerting both antiepileptic and pro-epileptic effects. The cleavage of its main receptor, full-length tropomyosin-related kinase B (TrkB-FL), was suggested to occur in status epilepticus (SE) in vitro. Moreover, under excitotoxic conditions, TrkB-FL was found to be cleaved, resulting in the formation of a new intracellular fragment, TrkB-ICD. Thus, we hypothesized that TrkB-FL cleavage and TrkB-ICD formation could represent an uncovered mechanism in epilepsy.

We used a rat model of mesial temporal lobe epilepsy (mTLE) induced by kainic acid (KA) to investigate TrkB-FL cleavage and TrkB-ICD formation during SE and established epilepsy (EE). Animals treated with 10 mg/kg of KA exhibited TrkB-FL cleavage during SE, with hippocampal levels of TrkB-FL and TrkB-ICD correlating with seizure severity. Notably, TrkB-FL cleavage and TrkB-ICD formation were also detected in animals with EE, which exhibited spontaneous recurrent convulsive seizures, neuronal death, mossy fiber sprouting, and long-term memory impairment. Importantly, hippocampal samples from patients with refractory epilepsy also showed TrkB-FL cleavage with increased TrkB-ICD levels. Additionally, overexpression of TrkB-ICD in the hippocampus of healthy rodents resulted in long-term memory impairment.

Our findings suggest that TrkB-FL cleavage and the subsequent TrkB-ICD production occur throughout epileptogenesis, with the extent of cleavage correlating positively with seizure occurrence. Moreover, we found that TrkB-ICD impairs memory. This work uncovers a novel mechanism in epileptogenesis that could serve as a potential therapeutic target in mTLE, with implications for preserving cognitive function.

## Introduction

Brain-derived neurotrophic factor (BDNF) is a neurotrophin that, through the activation of its high-affinity full-length tropomyosin-related kinase B receptor (TrkB-FL), promotes important biological actions such as neuronal survival, differentiation, growth, synaptic plasticity, and neurogenesis (1,2). Changes in BDNF signaling have been implicated in several pathological conditions, such as epilepsy. Indeed, both BDNF and TrkB-FL were reported to be increased in the hippocampus of patients with intractable temporal lobe epilepsy (TLE) (3,4). Interestingly, in in vivo animal models of epilepsy, BDNF levels were also shown to be upregulated (5,6). However, the in vivo data on TrkB-FL levels remain contradictory. TrkB-FL protein levels were described to be decreased after status epilepticus (SE) (7–9) and increased during established epilepsy (EE) (10), whereas TrkB-FL mRNA levels were shown to be unchanged (5,6) or increased (11,12).

Intriguingly, in experimental models, BDNF has been reported to exhibit both pro-epileptic and antiepileptic actions (13,14). In particular, it was shown that during acute seizures, BDNF levels are upregulated and TrkB-FL is activated, contributing to hyperexcitability and inducing more seizures in a positive feedback loop (13,14). Contrarily, chronic administration of BDNF was shown to decrease seizure frequency in an animal model of epilepsy (15,16).

The integrity of the TrkB-FL receptor is crucial for BDNF-mediated signaling. Under conditions of excessive glutamate release, such as in Alzheimer’s disease (AD), calpains are activated, leading to TrkB-FL cleavage and the consequent loss of BDNF signaling (8,17–19). This cleavage originates two protein fragments: a new membrane-bound truncated receptor (TrkB-T’) and an intracellular fragment (TrkB-ICD) (18). Truncated forms of TrkB-FL are dominant negative modulators of TrkB-FL and have been linked to neurodegeneration as a result of an imbalance in the TrkB-FL/truncated receptors ratio (8,17,20,21). In SE models, using hippocampal neurons exposed to MgCl_2_-free medium (21) or intrahippocampal administration of kainic acid (KA) (8), TrkB-FL levels were found to decrease while the levels of the naturally truncated TrkB isoforms increased. However, TrkB-ICD levels were not assessed in these studies. TrkB-ICD, with a half-life of 8 hours in neurons, can be localized in the cytosol and nucleus, and due to its tyrosine kinase activity, it can induce protein phosphorylation (22). Notably, our recent findings demonstrated that TrkB-ICD hyperpolarizes the resting membrane potential, increases the frequency of miniature excitatory postsynaptic currents (mEPSCs), and induces changes in the transcriptome, negatively impacting synaptic plasticity (23).

Since seizures pose an excitotoxic challenge and TrkB-ICD overexpression affects excitability and synaptic plasticity, we hypothesized that seizures could induce TrkB-FL cleavage, with TrkB-ICD formation potentially contributing to memory impairment. In this study, we aimed to investigate whether TrkB-FL cleavage occurs in a rat model of mesial temporal lobe epilepsy (mTLE) induced by KA at two timepoints: during SE and in EE. TrkB-FL cleavage was also studied in human hippocampal samples from patients with refractory epilepsy. Furthermore, TrkB-ICD was overexpressed in healthy rodents to investigate its effects on memory.

## Materials and Methods

### Animals

Male Sprague-Dawley rats (4 and 5 weeks old) and male C57BL/6J mice (12 weeks old) were purchased from Charles River Laboratories (Barcelona, Spain). The animals were housed in a virus antibody-free rodent facility, except those undergoing surgical procedures, which were maintained in a specific pathogen-free facility. Rats were housed in groups of four in individually ventilated cages enriched with corncob bedding, nesting materials, red acrylic tubes, and wooden blocks (all from LBS Biotechnology). Mice were housed in groups of four/five in individually ventilated cages enriched with corncob bedding, cardboard tubes, and nesting materials (from LBS Biotechnology). Animals received food (autoclaved diet pellets) and water (sterile water treated by reverse osmosis) *ad libitum*. Temperature (20-24°C), humidity (55% ± 10%), and light/dark cycle (14 h light:10 h dark; lights on between 7 a.m. and 9 p.m.) of the facilities were controlled. All experiments were performed between 8 a.m. and 7 p.m.

After one week of acclimatization in the rodent facility, each animal was handled for five minutes, every day, for five consecutive days to be familiarized with the experimenter and reduce stress during procedures. Animals subjected to stereotaxic surgery were handled before behavioral tests. Animals were handled according to standard and humanitarian procedures to reduce animal suffering. All procedures met the European Community guidelines (86/609/EEC; 2010/63/EU; 2012/707/EU) and Portuguese legislation (DL 113/2013) for the protection of animals used for scientific purposes. The protocol was approved by the institutional animal welfare body, ORBEA-iMM, and the competent national authority, *Direcção Geral de Alimentação e Veterinária* (DGAV), under the project reference 002477\2021.

### Experimental model of mesial temporal lobe epilepsy induced by kainic acid administration

Different dose regimens of KA were tested to optimize a protocol to generate an mTLE model with a low mortality rate, with at least 50% of the animals presenting stage 5 seizures during SE, and stage 4/5 spontaneous seizures in EE. To achieve this, 7-week-old male Sprague-Dawley rats (250 g) were treated with a single intraperitoneal (i.p.) injection of KA (Tocris, Bristol) at several doses (5 mg/kg, 8.5 mg/kg, or 10 mg/kg) or three injections of KA repetitive low dose (5 mg/kg) with a 30-minute interval between each injection. Antiseizure drugs were not administered to terminate SE. Control animals were injected with the vehicle solution (0.9% NaCl). The animals were assessed at two different periods: during SE (Figure 1a) or in EE (Figure 1b).

**Figure 1.**
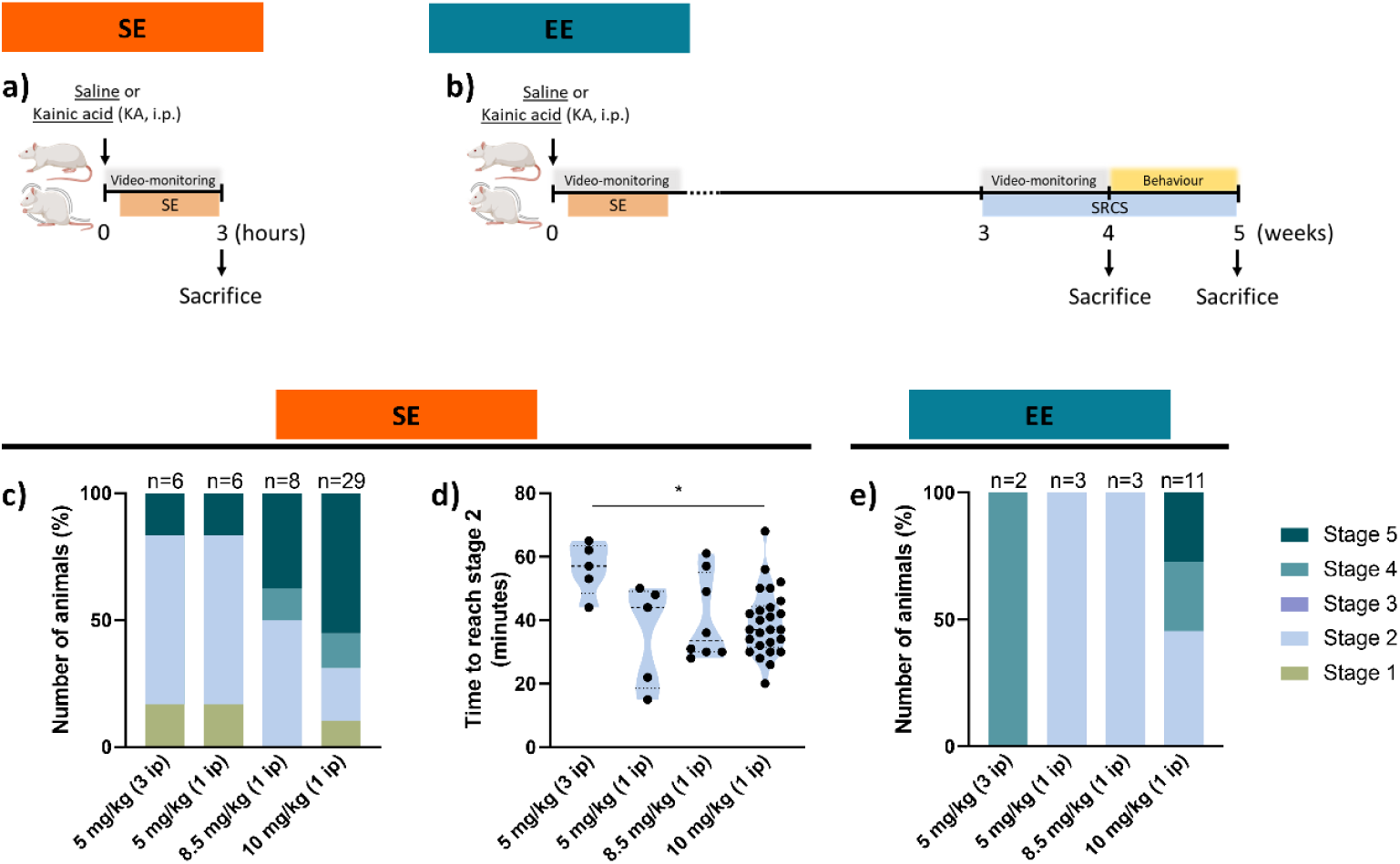
Experimental design of kainic acid (KA)-induced model of mesial temporal lobe epilepsy (mTLE) during status epilepticus (SE) (a) or during established epilepsy (EE) (b). (a) Animals were sacrificed 3 hours after the KA injection during SE. (b) Animals were sacrificed during EE at 4 weeks after the KA injection or 5 weeks after the KA injection after performing behavioral tests. (c-e) The seizure stage during SE and during EE depends on the dose of KA. (c) Distribution of maximum seizure stage reached during SE after different intraperitoneal (i.p.) dose regimens of kainic acid (KA). (d) Time to reach seizure stage 2 after different dose regimens of KA (n=5-26). (e) Distribution of maximum seizure stage reached during EE, after SE induction by different dose regimens of KA. Statistical analysis was performed using Brown-Forsythe ANOVA and Welch ANOVA tests followed by Dunnett’s T3 multiple comparisons test. i.p., intraperitoneal injection. SRCS, spontaneous recurrent convulsive seizures. Panels a and b created with BioRender.com.

Out of the initial cohort of 49 animals injected with KA, 29 [5 mg/kg (3 i.p.): n=4; 5 mg/kg (1 i.p.): n=3; 8.5 mg/kg (1 i.p.): n=5; 10 mg/kg (1 i.p.): n=17] were sacrificed at 3 hours after SE induction, corresponding to the SE group (Figure 1a). The other 19 animals [5 mg/kg (3 i.p.): n=2; 5 mg/kg (1 i.p.): n=3; 8.5 mg/ kg (1 i.p.): n=3; 10 mg/ kg (1 i.p.): n=11] were sacrificed to constitute the group with EE (Figure 1b). All animals from this group were sacrificed 4 weeks after SE induction, except 5 animals treated with 10 mg/kg of KA used in behavioral tests and sacrificed 5 weeks after SE induction. Only one animal treated with 10 mg/kg of KA (not included in the previous numbers) died the day after SE and was only included in the SE seizure stage assessment. Detailed information per animal and its behavioral and molecular assessments can be found in Supplementary Table 1 and Supplementary Table 2.

During the KA injection day, animals were closely monitored by an experienced experimenter, and video recorded using two cameras located on opposite sides of the cage (Go pro 4K and USB webcam-Model: ELP-USBFHD05MT-DL170). The video recordings were used to assess seizure severity. Seizures were classified according to the modified Racine Scale: stage 1, immobility and facial clonus; stage 2, nodding and wet-dog shaking; stage 3, unilateral forelimb clonus with lordotic posture; stage 4, lateral forelimb clonus with rearing; and stage 5, bilateral forelimb clonus with rearing and falling (24,25). According to the pre-defined humane endpoint, animals presenting stage 5 seizures for more than five consecutive minutes were sacrificed.

Animals of the EE group were monitored for the appearance of distress symptoms in the following days. In the fourth week after SE induction, the animals were again video recorded between 6 p.m. and 9 a.m. for 4 consecutive nights using an ELP camera and SplitCam v 10.5.12 software to evaluate the appearance of spontaneous recurrent convulsive seizures (SRCS). All animals were sacrificed with a lethal dose (1-2 mL) of sodium pentobarbital (Euthasol®, 400 mg/ml, Dechra).

### Seizure severity assessment

The assessment of seizure severity was conducted retrospectively by analyzing the video recordings captured during the SE day and at the four-week timepoint following KA administration.

During SE, seizures followed a progressive pattern from stage 1 to stage 3 and then occurred interchangeably between stages 3 and 5 (stages of convulsive seizures). Before the onset of seizures, animals displayed specific features such as ear movements and mouth opening and closing. Foaming at the mouth was often observed during seizures in stages 4-5. Animals presenting at least two wet-dog shakes were considered in stage 2. A convulsive seizure was only considered terminated when the animal placed both paws on the ground for at least 2 seconds. At the end of SE, animals primarily displayed stage 3 seizures or head automatisms. Approximately 4 hours after the KA injection, animals stopped displaying convulsive seizures and started a latent period exhibiting their typical behaviors such as walking or eating. Four weeks after SE induction some animals displayed SRCS after the latent period.

During SE, the following seizure parameters were measured: time to reach stage 2 seizures; maximum seizure stage; number of seizures, and SE duration (time between the first stage 3 seizure and animal sacrifice). During EE, the maximum seizure stage and the number of seizures per hour were quantified. Seizures during SE and EE were assessed by the same experimenter.

### Experimental design for TrkB-ICD overexpression experiments

TrkB-ICD overexpression was induced in both mice and rats, with animals randomly allocated to the different experimental groups.

Mice (28 g) were injected with lentiviral vectors at 12 weeks of age, allowing lentiviral expression for 6 weeks before behavioral assessment. Behavioral tests were conducted over 2 weeks after which the mice were sacrificed for electrophysiological studies. A total of 51 mice (CTL: 15; PBS: 10; eGFP: 12; TrkB-ICD: 14) were used in the behavioral experiments. A subset of 24 mice from the initial pool of 51 mice (CTL: 6; PBS: 5; eGFP: 7; TrkB-ICD: 6) was used for the electrophysiological experiments.

Rats were injected with lentiviral vectors at 5 weeks of age, with lentivirus expression allowed for 5 weeks before behavioral assessment. Behavioral tests were performed over 1 week after which rats were sacrificed for immunohistochemistry analysis. A total of 15 rats (eGFP: 7; TrkB-ICD: 8) were included in the behavioral experiments.

### Stereotaxic surgery for in vivo lentiviral transduction

Mice were anesthetized with 2-3 % isoflurane (IsoFlo® 1000 mg/g, Esteve) in 100 % oxygen delivered through a nozzle mask at a continuous flow rate of 1-2 mL/min. Similarly, rats were anesthetized with 3-3.5 % isoflurane at a continuous flow rate of 3-3.5 mL/min, or by administering an i.p. injection of an anesthetic solution containing 75 mg/kg ketamine (Ketamidor® 100 mg/ml; Richter Pharma) and 1 mg/kg medetomidine (Domtor® 1mg/ml; Ecuphar). To reverse the i.p. anesthesia in rats, atipamezole (Antisedan® 5 mg/ml; Ecuphar) was administered at 1mg/kg per body weight.

Animals were maintained above heating pads throughout the procedure to prevent hypothermia. To minimize animal discomfort the following agents were administered: buprenorphine (Bupaq® 0.3 mg/mL; Richer Pharma) for analgesia, Iodopovidone (Betadine® 100 mg/mL, Meda pharma) as an antiseptic, ophthalmic gel (Lubrithal, Dechra) to protect the eyes from dryness, and lidocaine (Lidonostrum® 20 mg/g, Sidefarma) as a local anesthetic for the ear canal. Animals were positioned in a stereotaxic frame (Stoelting, adapted for each rodent) in a flat-skull position.

The protein of interest, TrkB-ICD, and the control protein, eGFP, were overexpressed using pFCK-CamKII-TrkB-ICD-ZsGreen or pFCK-CamKII-eGFP lentiviral vectors, respectively. These lentiviral vectors previously used (23) were diluted in sterile phosphate-buffered saline (PBS; 137 mM NaCl, 2.7 mM KCl, 1.8 mM KH_2_PO_4_, and 10 mM Na_2_HPO_4_.2H_2_O dissolved in *Mili*-Q® water, pH 7.40) the day before, and then injected using a Hamilton 25 μL syringe (702 N, ga 29/51mm, PST3, Hamilton Company). A total of 3×10^7^ Transduction Units constituted the lentiviral solution. Each solution (mice: 1 μL, rats: 2 μL; per injection site) was bilaterally injected into the hippocampi (mice: CA1: AP -1.9, DV -1.4, ML ± 1.5; CA3: AP -1.9, DV -2.2, ML ± 2.3.

Rats: CA1: AP -3.6, DV -2.5, ML ± 2; CA3: AP -3.6, DV -3.8, ML ± 4.1) at a rate of 400 nL/min, totalizing four injections per animal. The needle remained in place (mice: 1 minute, rats: 5 minutes) before being slowly raised at a 0.5 mm/min rate. Following the four intrahippocampal injections, animals were sutured, and Dexpanthenol (Bepanthene® 50 mg/g: Bayer) was applied. Animals were placed in an isolated cage with special bedding until recovery from the anesthesia and then returned to the original cage.

### Behavioral tests

*mTLE experiments*: In the fifth week after KA or saline injection, rats performed the following behavioral tests: elevated plus maze (EPM), open field (OF), and object location test.

*TrkB-ICD overexpression experiments*: In the sixth week after lentiviral injection, mice were subjected to the following behavioral tests: EPM, OF, novel object recognition (NOR) test, and Morris water maze (MWM) test. In the fifth week after lentiviral injection, rats performed the following behavioral tests: EPM, OF, and object location test.

In all tests, each rodent had an arena appropriate for its size. The maze/arena and objects were cleaned with 30% ethanol solution between animals to erase any olfactory clues. Dim light was used. All tests were video-recorded, and animals were live-tracked using the video-tracking software SMART© VERSION 3.0.06 (EXTENSION GA, TW).

#### Elevated plus maze test

Anxiety-like behavior was assessed in the EPM test, as previously described (26,27). A plus sign-shaped maze with two open (mice: 5×29 cm, rats: 50×10 cm) and two closed arms (mice: 5×29×15 cm, rats: 50×10×50 cm), elevated above the floor (mice: 50 cm, rats: 65 cm), was used (Figure 5f-*rat apparatus*). Each animal was placed in the center of the maze, with the head facing an open arm, and allowed to explore for 5 minutes.

During acquisition, virtual areas (for each arm and the center) were designed to measure the different parameters (time spent in the open arms, number of entries in the open arms, and the total distance traveled) using the software.

For the analysis, the criterion for transitioning between zones was based on the center of mass of the animal.

#### Open field test

Animal locomotor activity was measured in the OF test, as previously described (27). Each animal was positioned in the center of an empty square box (mice: 40×40×40 cm, rats: 60×60×40 cm) and allowed to explore (mice:15 minutes; rats: 10 minutes) for 3 consecutive days. This constitutes the habituation phase of the object location test and the NOR test. The OF test was evaluated in the first 5 minutes of the first day. During acquisition, three virtual areas (Figure 5g-*rat apparatus*) were designed to measure the different parameters (time spent in the center zone, total distance traveled, and mean speed in the center zone) using the software. For the analysis, the criterion for transitioning between zones was based on the center of mass of the animal.

#### Novel-object recognition test

The NOR test was used to evaluate long-term memory in mice, as previously described (26). After the habituation phase (3 days in the OF arena), two identical objects (familiar objects: A and A’) were placed inside the arena and mice were allowed to freely explore for 5 minutes (training day). After a 24-hour retention interval, one of the familiar objects was swapped for a novel object (B) and animals were allowed to freely explore for 5 minutes (test day). Several aspects were carefully planned to ensure an unbiased study: objects used on both days (wooden dolls, 7×6 cm) were optimized for mice; object location was randomized for all animals; object role (novel and familiar) was randomized inside the same experimental group; both objects were used as the novel object the same number of times for all experimental groups; the objects were attached to the bottom of the arena with velcro, unseen to the animals).

The investigator performed a blind analysis for all experimental groups to assess the time each animal spent exploring each object. Exploration time (in seconds) was considered when the animal spent time close (<1 cm) to the objects, including smelling the object, rearing up towards the object, or touching it. Importantly, animals that failed to explore either object for more than 10 seconds were excluded. The percentage of exploration time was given by the time spent exploring an object divided by the time spent exploring both objects (in seconds) and multiplied by 100. The novelty preference index (NPI) was calculated by subtracting the exploration time of the familiar object from the exploration time of the novel object and then dividing the result by the total exploration time of both objects. The NPI ranges from -1 to 1, where 1 indicates exclusive exploration of the novel object, 0 indicates equal exploration time for both objects, and -1 indicates no exploration of the novel object.

#### Object location test

The object location test was used to evaluate hippocampal-dependent long-term memory in rats (28). After the habituation phase (3 days in the OF arena), each rat was exposed for 5 minutes to two equal objects (green octahedrons, Figure 5h), which were placed in opposite corners of the same wall of the arena (training day). Each animal was placed into the arena as far away from the objects as possible, facing away from the objects, to avoid introducing any object bias. After 24h, one of the objects was moved to a different location and the other object was retained in the same position as in the habituation phase (Figure 5h), and each animal was allowed to explore both objects for 3 minutes (test day).

The displaced object and its position inside the arena were counterbalanced among animals. Exploration was considered whenever the head of the animal was inside a virtual circle of 5 cm delimited around each object. The percentage of exploration ratio was given by the time spent exploring an object divided by the time spent exploring both objects (in seconds) and multiplied by 100. For the analysis, the criterion for transitioning between zones was based on the head of the animal.

#### Morris water maze test

The MWM test was performed to assess hippocampal-dependent learning and long-term spatial memory in mice, as previously described (23). The test was executed for 5 consecutive days. On the first day, the habituation day, each animal was familiarized with the presence of an escape platform, placed in the center of the tank, 1 cm above water level. In the following 3 days, the acquisition phase, each animal had four swimming trials (60 seconds) per day (with 30-minute intervals between trials) to find the escape platform submerged 1 cm below water level, in a fixed position. The platform was randomized for each animal, in the center of the target quadrant. Finally, on the fifth day, the probe test, the escape platform was removed, and animals were allowed to swim freely for 60 seconds. The time each mouse spent in the target virtual quadrant, where the platform was previously located, as well as the number of entries into that quadrant, were measured. For the analysis, the criterion for transitioning between zones was based on the center of mass of the animal.

### Perfusion and tissue processing

Mice, under deep isoflurane anesthesia, were sacrificed by decapitation. Then, the brain was removed from the skull and placed on ice-cold artificial cerebrospinal fluid (aCSF; 124 mM NaCl, 3 mM KCl, 1.2 mM NaH_2_PO_4_.H_2_O, 25 mM NaHCO_3_, 10 mM glucose monohydrate, 2 mM CaCl_2_ and 1 mM MgSO_4_, dissolved in *Mili*-Q® water, pH 7.4). The hippocampi were isolated, and 400 μm thick coronal sections were cut using a McIlwain tissue chopper (Campden Instruments). The sections were placed in a petri dish filled with aCSF and observed under a widefield fluorescence microscope (Axiovert 200, ZEISS) with the AxioVision 4 software (ZEISS). Only hippocampal sections displaying green fluorescence from eGFP or ZsGreen probes were selected for electrophysiological experiments.

Rats were euthanized with a lethal dose of sodium pentobarbital, followed by transcardial perfusion with PBS, for 20 minutes. After perfusion, the rat was decapitated and the brain was removed and placed on ice-cold aCSF. The two hemispheres were separated, and the hippocampus and whole cortex from the right hemisphere were dissected and stored at -80 °C until further use for western blotting. The left hemisphere was postfixed with 4% paraformaldehyde (PFA) (dissolved in *Mili*-Q® water, pH 6.8-7.4) for 72 hours at 4 °C, followed by cryoprotection in sequential 15% and 30% sucrose in PBS (4 °C, until the tissue sinks). Later, the hemisphere was gelatine-embedded and coronally sectioned between anteroposterior coordinates -2 mm and -4.8 mm (dorsal hippocampus) with a cryostat (Leica CM3050 S, Leica Biosystems). Sections of 40 µM were placed on anti-freeze solution [30% glycerol, 30% ethylene glycol, 0.1 M phosphate buffer (0.5 M Na_2_HPO_4_.2H_2_O adjusting pH 7.3-7.4 with 0.5 M NaH_2_PO_4_.2H_2_O, both dissolved in *Mili*-Q® water) diluted in *Mili*-Q® water] and stored at -20 °C until further use for immunohistochemistry analyses.

### Human samples

The use of human samples collected from patients with epilepsy was approved by the Ethics Committee from *Centro Hospitalar Universitário Lisboa Norte* and *Centro Académico de Medicina de Lisboa* (n° 207/21) and met the Portuguese Law on Clinical Research (Law No. 21/2014, 16 April), amended in law (Law No. 73/2015, 27 July). Biobank from *Instituto de Medicina Molecular João Lobo Antunes* provided frozen samples of the hippocampus from 11 patients who underwent temporal lobe resections due to refractory epilepsy, after an extensive presurgical evaluation at *Centro de Referência para a área da epilepsia refratária*, member of the ERN EpiCARE for Complex and rare epilepsies (details in Supplementary Table 3).

Additionally, post-mortem hippocampal samples (n=7) included in this study (details in Supplementary Table 4), were obtained from the archives of the Department of Neuro(Pathology) of the Amsterdam University Medical Centers (Amsterdam UMC-location AMC, the Netherlands). These samples were collected from autopsies of patients with no history of neurological diseases. Tissue was obtained and used following the Declaration of Helsinki, the Amsterdam UMC research code, and a protocol approved by the local Biobank Research Ethics Committee. All samples were maintained at -80 °C until further processing for western blotting.

### Western blotting

Hippocampal (human or rat) and cortical (rat) samples were lysed in 1.2 mL of radioimmunoprecipitation assay (RIPA) lysis buffer composed of 4% NP-40, 40 mM Tris-HCl, 1 mM EDTA, 150 mM NaCl and 0.1% SDS 10%, 10 mM NaF, 5 mM Na_3_VO_4_, and protease inhibitors (complete^TM^ Mini EDTA-free Protease Inhibitor Cocktail, Roche), dissolved in *Mili*-Q® water. After homogenization using a sonicator (Soniprep MSS 150.CX3.5: Sanyo), total protein was quantified using the Bio-Rad *DC* Protein Assay Kit (Bio-Rad), and absorbance was read at 750 nm (Infinite M200, Tecan Trading AG). Lysates were mixed with 6x sample buffer (12% SDS, 0.06% bromophenol blue, 47% glycerol, 600 mM dithiothreitol, 60 mM Tris-HCl, dissolved in *Mili*-Q® water, pH 6.8) and denatured for 10 minutes, at 95 °C. Each sample (40 μg total protein/well) and 5 μl of molecular marker (NZYColour Protein Marker II: Nzytech) were resolved on 10% SDS-polyacrylamide (SDS-PAGE) gel within a running buffer (25 mM Tris-base, 192 mM glycine, 10 ml SDS 10%, dissolved in *Mili*-Q® water, pH 8.3) at 80 volts, until the marker started to separate and then, at 120 volts for approximately 90 minutes using Mini-Protean® Tetra System (Bio-Rad). SDS-PAGE-separated proteins were transferred to polyvinylidene fluoride (PVDF) membrane (Immun-Blot® PVDF Membranes for Protein Blotting, Bio-Rad) with a transfer buffer (25 mM Tris-base, 192 mM glycine, 10% methanol, dissolved in *Mili*-Q® water, pH 8.3) using the same system at 350 mA for 105 minutes. Membranes were blocked in 3% bovine serum albumin (BSA) in TBS-T (20 mM Tris-base, 137 mM NaCl, 0.1% Tween-20, dissolved in *Mili*-Q® water, pH 7.6) for 1 hour at room temperature (RT). Membranes were incubated on a shaker with the primary antibodies: rabbit polyclonal anti-Trk(C14) (1:1 000, Santa Cruz Biotechnology, #sc-11, RRID: AB_632554), mouse monoclonal anti-Glyceraldehyde-3-Phosphate Dehydrogenase (GAPDH, 1:5000, Thermofisher, #AM4300, RRID: AB_2536381), overnight, at 4 °C. GAPDH was used as internal control. Then membranes were incubated with horseradish peroxidase (HRP)-conjugated secondary antibodies (goat anti-rabbit IgG-HRP, 1:10 000, Bio-Rad, # 170-6515, RRID: AB_11125142; goat anti-mouse IgG-HRP, 1:10 000, Bio-Rad, # 170-6516, RRID: AB_11125547), for 1 hour at RT. All antibodies were diluted in 3% BSA in TBS-T. The membranes were rinsed in TBS-T (3 times x 5 minutes) between every step of the protocol.

Chemiluminescence signals were developed using enhanced chemiluminescence detection (Western Lightning® ECL Pro: PerkinElmer Inc) and detected by the ChemiDoc XRS+ System (Bio-rad) or Amersham ImageQuant 680 system (Cytvia). Images were developed with a total of 50 pictures and a cumulative exposure time per image ranging between 10 to 20 seconds. Band intensities were quantified by digital densitometry using Fiji (National Institutes of Health), maintaining the same area between bands within the same membrane. The intensity of each protein band was normalized to the band intensity of GAPDH and then to the mean of controls within the same membrane.

### Immunohistochemistry

Gelatin was removed by sequentially immersing free-floating slices in PBS at 37 °C. Slices were then incubated with blocking and permeabilization solution (3% BSA, 0.2% Triton in PBS) for 1 hour at RT. Particularly, slices from animals injected with lentiviral vectors were incubated with 0.1 M glycine in PBS for 20 minutes after the blocking solution. Later, the primary antibodies [rabbit monoclonal anti-NeuN (D3S3I) (1:500, Cell Signaling Technology, #12943, RRID: AB_2630395), rabbit polyclonal anti-Synaptoporin (1:500, Synaptic Systems, #102 002, RRID: AB_887841), rabbit monoclonal anti-Pan-Trk [EPR17341] (1:500, Abcam, #ab181560, RRID: AB_2940902), mouse monoclonal anti-TrkB (Clone 47/TrkB) (1:500, BD Transduction Laboratories, #610101, RRID: AB_397507)] diluted in the blocking and permeabilization solution were incubated for 2 overnights at 4°C. Subsequently, slices were incubated with the secondary antibodies [donkey anti-rabbit IgG (H+L) Alexa Fluor™ 568 (1:500, #A10042, RRID: AB_2534017), donkey anti-rabbit IgG (H+L), Alexa Fluor™ 647 (1:500, #A31573, RRID: AB_2536183), donkey anti-mouse IgG (H+L), Alexa Fluor™ 568 (1:500, #A10037, RRID: AB_2534013), all from Thermo Fisher Scientific] and 4′,6-diamidino-2-phenylindole (DAPI, 1:1000, Sigma-Aldrich, #D9564) diluted in PBS for 105 minutes. Slices were mounted on microscope slides (1 mm SuperFrost Plus adhesion slides), with a mowiol fluorescent medium (2.4 g Polyvinyl alcohol 4-88, 6.1 g glycerol, 12 mL Tris-base (200 mM, pH 8.0), and 6 mL *Mili*-Q® water), covered with high-precision cover glasses (thickness No. 1.5H), and left to dry in the dark for 2 days. Images were captured in the entire thickness (z-stack) of the slice on a Cell Observer SD (ZEISS) with an LD C-Apochromat 40x/1.1 water Correction M27 objective. Additionally, a few images of slices from animals injected with lentiviral vectors were captured in Confocal 980 (ZEISS) with a Plan-Apochromat 63x/1.4 Oil DIC M27. ZEN 3.2 (blue edition, ZEISS) software was used for the analysis.

### Extracellular ex vivo recordings

Slices displaying fluorescence were allowed to recover functionally and energetically for 1 h in a resting chamber, filled with aCSF, at RT. Then, slices were transferred to a submerging chamber and continuously superfused at a 3 mL/min rate with gassed aCSF at 32 °C. A microelectrode filled with 4 M NaCl (2–8 MΩ resistance) placed in the *stratum radiatum* of the CA1 area was used to record extracellularly evoked field excitatory postsynaptic potentials (fEPSP). A bipolar concentric wire electrode positioned on the Schaffer collateral/ commissural fibers was used to stimulate, at 200 μA, this pathway (rectangular pulses of 0.1 milliseconds duration), every 20 seconds. The slope of the initial phase of the potential was quantified, and averages of six consecutive fEPSP were obtained and recorded using an Axoclamp 2B amplifier (Axon Instruments), digitized and continuously stored on a computer with the WinLTP software (WinLTP Ltd. and The University of Bristol)(29).

#### Long-term potentiation induction

Long-term potentiation (LTP) was induced, after a stable period of at least 10 minutes, by θ-burst protocol consisting of 3 trains of 100 Hz, 3 stimuli, separated by 200 ms (30). This θ-burst protocol was chosen as it mimics the physiological stimulation pattern observed in the hippocampus of live animals during learning and memory episodes (31). The stimulus intensity remained constant. LTP magnitude was quantified as the percentage of change in the average slope of the fEPSP 50 to 60 min after LTP induction, relative to the average slope of the fEPSP measured during the 10 min that preceded the induction.

#### Post-tetanic potentiation

Post-tetanic potentiation (PTP) was quantified by calculating the average of the first three slope responses following LTP induction.

### Statistical analyses

Before statistical analysis, data normality was assessed via the Shapiro-Wilk test. For data not following a normal distribution, a logarithmic transformation (Y=Log(Y)) was applied and the normal distribution was reevaluated.

For data following a normal distribution and assuming all groups were from populations with equal variances (similar (±0.01) standard deviation), unpaired *t*-tests were implied to compare the means between two groups, or a one-way ANOVA to compare among multiple groups. ANOVA was followed by Dunnett’s test for multiple comparisons.

For data following a normal distribution and not assuming all groups were from populations with equal variances (different standard deviations), unpaired *t*-tests with Welch’s correction were used to compare the means between two groups, or a Brown-Forsythe and Welch ANOVA tests to compare among multiple groups. Brown-Forsythe and Welch ANOVA tests were followed by Dunnett’s test for multiple comparisons.

In the case of non-normality, Mann-Whitney U tests were used to compare the ranks between two groups, or Kruskal-Wallis tests to compare among multiple groups. Kruskal-Wallis tests were followed by Dunnett’s test for multiple comparisons.

Paired t-test was also used to compare the same animal at two different timepoints, and mixed-effects analysis was used to compare multiple groups at multiple timepoints. One animal from the EE group was an outlier by the Grubb’s method (alpha=0.05) in the TrkB-ICD protein levels and was excluded from all western blot analyses. Using the same statistical method, one mouse overexpressing eGFP was excluded from the novel object recognition test since it was an outlier during the training day.

A correlation was performed between different variables (seizure stage, number of seizures, and the protein levels of TrkB-FL, TrkB-ICD, and TrkB-ICD/TrkB-FL, either during SE or EE). Pearson correlation was used when the data followed a normal distribution, and nonparametric Spearman correlation implied when data did not follow a normal distribution.

All statistical analyses were performed using GraphPad Prism version 8.0.0 for Windows (GraphPad Software). Statistical significance was set at *p* < 0.05 and a statistical tendency was considered when 0.05 < *p* < 0.1.

## Results

### Seizure severity depends on the dose of KA

To establish an mTLE rat model, several KA dose regimens were tested to select the appropriate conditions. These included 3 doses of 5mg/kg; a single dose of 5 mg/kg; a single dose of 8.5 mg/kg and a single dose of 10 mg/kg (Supplementary Table 1 and 2). To assess the seizure stage during SE using the modified Racine scale, all animals were grouped regardless of whether they were sacrificed 3 hours or 4-5 weeks after SE induction. The administration of 5 mg/kg of KA, in a single dose or in three doses, yielded similar outcomes during SE: 16.7% of the rats remained in stage 1, 66.7% experienced stage 2 seizures, while only 16.7% reached stage 5 seizures (Figure 1c). With 8.5 mg/kg of KA, 50% of the animals exhibited stage 2 seizures, 12.5% experienced stage 4 seizures, and 37.5% reached stage 5 seizures (Figure 1c). Notably, administration of 10 mg/kg of KA resulted in stage 1 in 10.4% of the animals, stage 2 seizures for 20.7%, and stage 4 seizures for 13.8%, while 55.2% of animals reached stage 5 seizures (Figure 1c). In addition, animals treated with 10 mg/kg of KA experienced stage 2 seizures earlier than animals treated with multiple KA doses of 5 mg/kg (Figure 1d).

Four weeks after inducing SE, animals treated with a single dose of 5 mg/kg or 8.5 mg/kg of KA experienced only stage 2 seizures and did not progress to SRCS (stages 3-5) during EE (Figure 1e). All animals treated with three doses of 5 mg/kg depicted stage 4 seizures, while 54.5% of the animals treated with 10 mg/kg of KA showed spontaneous stage 4 or 5 seizures (Figure 1e).

Therefore, the administration of 10 mg/kg of KA induced stage 5 seizures during SE and led to the development of SRCS in more than 50% of the animals during EE.

### TrkB-FL and TrkB-ICD protein levels are altered depending on the KA-administered dose

To evaluate the putative TrkB-FL cleavage in the hippocampus and cortex during SE and EE, the levels of both TrkB-FL and its cleaved fragment, TrkB-ICD, were investigated by western blot (Figure 2, Supplementary Figure 1, Supplementary Figure 2).

**Figure 2.**
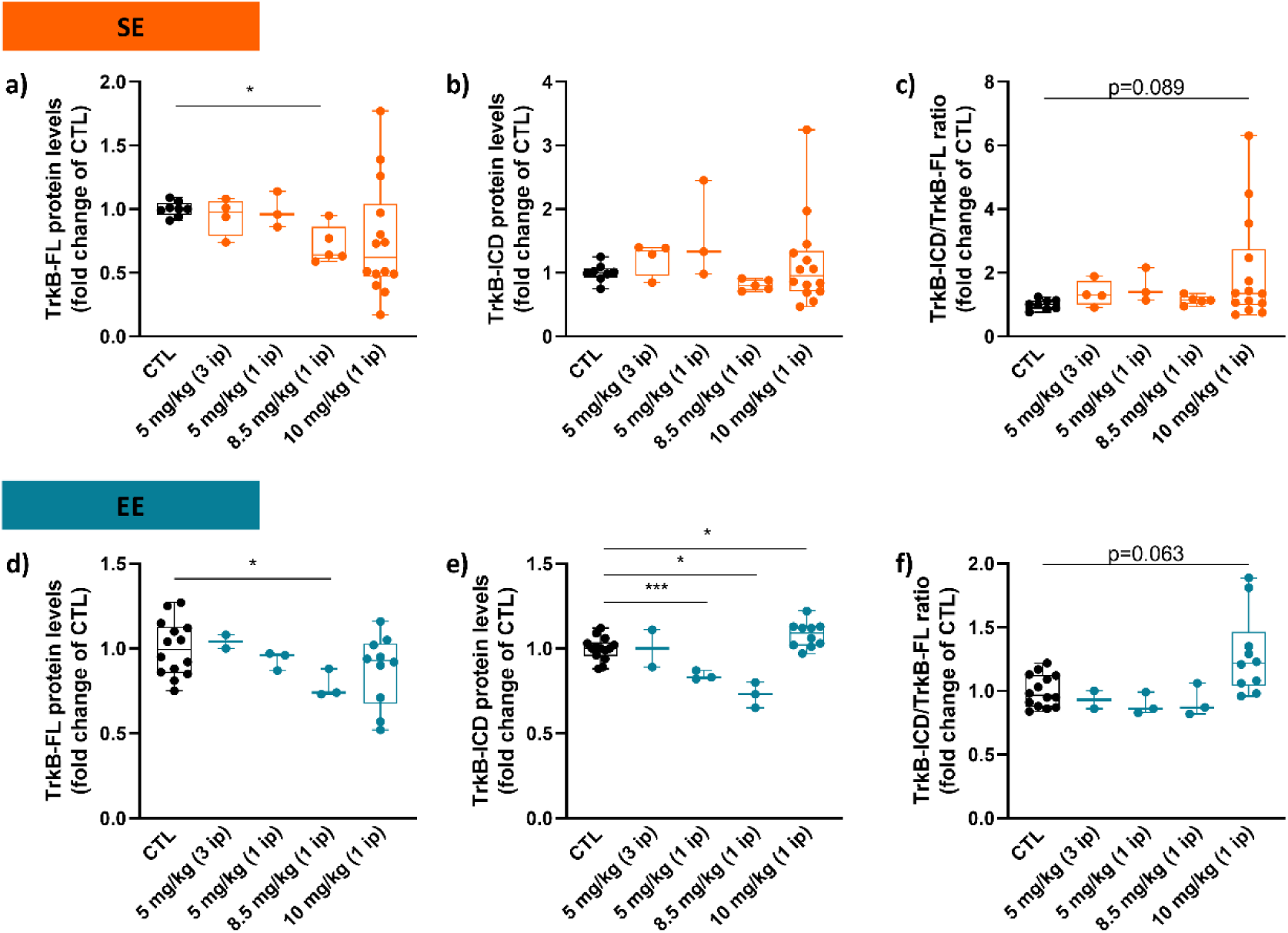
Hippocampal protein levels of animals treated with different dose regimens of kainic acid (KA) and sacrificed during status epilepticus (SE) (a-c, orange) or during established epilepsy (EE) (d-f, blue) are changed depending on the KA administered dose.(a) TrkB-FL levels, (b) TrkB-ICD levels, and (c) TrkB-ICD/TrkB-FL ratio of animals sacrificed on the SE day (CTL: n=8; 5 mg/kg (3 i.p.): n=4; 5 mg/kg (1 i.p.): n=3; 8.5 mg/kg (1 i.p.): n=5; 10 mg/kg (1 i.p.): n=14). (d) TrkB-FL levels, (e) TrkB-ICD levels, and (f) TrkB-ICD/TrkB-FL ratio of animals sacrificed during EE (CTL: n=14; 5 mg/kg (3 i.p.): n=2; 5 mg/kg (1 i.p.): n=3; 8.5 mg/kg (1 i.p.): n=3; 10 mg/kg (1 i.p.): n=10). Control animals in all experiments were treated with saline (black). Statistical analysis was performed using Brown-Forsythe ANOVA and Welch ANOVA tests (a, c, d, e, f) or Kruskal-Wallis test (b), all followed by Dunnett’s multiple comparisons tests. Results are shown as median, minimum, 25% and 75% percentile, and maximum. *p<0.05, versus CTL group.

In SE, animals treated with 8.5 mg/kg of KA revealed a significant decrease in hippocampal TrkB-FL protein levels compared with control animals (p=0.031, Figure 2a, Supplementary Figure 2). No significant differences in the protein levels of TrkB-FL or TrkB-ICD were observed in the hippocampus of animals treated with other dose regimens of KA compared with control animals (Figure 2a and b, Supplementary Figure 2). Although the highest dose of KA (10 mg/kg) resulted in considerable variability in hippocampal levels of both TrkB-FL and TrkB-ICD (Figure 2a and b, Supplementary Figure 2), a tendency toward an increased TrkB-ICD/TrkB-FL ratio in the hippocampus compared with control animals was observed (p=0.089, Figure 2c, Supplementary Figure 2).

Regarding the protein levels of TrkB-FL in the cortex during SE, the pattern was less clear than in the hippocampus. A tendency for decreased cortical levels of TrkB-FL was found in animals treated with a single dose of 5 mg/kg of KA (p=0.075 vs control, Supplementary Figure 1a, Supplementary Figure 2), while a tendency for increased levels was found in animals treated with 8.5 mg/kg of KA (p=0.059 vs control, Supplementary Figure 1a, Supplementary Figure 2). No differences were found in cortical levels of TrkB-ICD or TrkB-ICD/TrkB-FL ratio during SE among the different KA dose regimens (Supplementary Figure 1b and c).

In EE, animals treated with 8.5 mg/kg of KA also exhibited a significant decrease in hippocampal TrkB-FL protein levels when compared with control animals (p=0.043, Figure 2d, Supplementary Figure 2). However, a decrease in TrkB-ICD protein levels was found in the hippocampus of these animals (p=0.024), as well as, in animals treated with a single dose of 5 mg/kg of KA when compared with control animals (p=0.0002, Figure 2e, Supplementary Figure 2). While animals treated with 10 mg/kg of KA displayed a significant increase in TrkB-ICD protein levels (p=0.040, Figure 2e, Supplementary Figure 2) and a tendency towards an increase in the TrkB-ICD/TrkB-FL ratio (p=0.063, Figure 2f, Supplementary Figure 2).

With regard to the cortical levels during EE, no differences were found in TrkB-FL protein levels between animals treated with KA and control animals (Supplementary Figure 1d, Supplementary Figure 2). However, animals treated with 8.5 mg/kg of KA demonstrated an increase in TrkB-ICD protein levels (p=0.024, Supplementary Figure 1e, Supplementary Figure 2) and in the TrkB-ICD/TrkB-FL ratio (p=0.007, Supplementary Figure 1f, Supplementary Figure 2).

These results show that the 8.5 mg/kg of KA induces a decrease in the TrkB-FL hippocampal levels in SE and EE, but also a decrease in the TrkB-ICD hippocampal levels in EE. While the highest dose of KA, 10 mg/kg, induced a noteworthy trend towards an increased TrkB-ICD/TrkB-FL ratio in the hippocampus during SE and EE, with a significant increase of TrkB-ICD formation in EE, pointing to the occurrence of TrkB-FL cleavage.

### TrkB-FL levels are lower in KA-treated animals (10 mg/kg) with higher seizure stage in SE but not during EE

The considerable variability in hippocampal protein levels of TrkB-FL and TrkB-ICD observed in animals treated with 10 mg/kg of KA, led us to hypothesize that this variability could be related to the differences in seizure severity.

In SE, there was a negative correlation between seizure stage and TrkB-FL protein levels in the hippocampus of animals treated with 10 mg/kg of KA, indicating that animals with higher seizure stage had lower levels of TrkB-FL (p=0.043, Figure 3a). Additionally, TrkB-ICD levels were positively correlated with the number of seizures, implying that animals with more seizures during SE had higher levels of TrkB-ICD (p=0.044, Figure 3b).

**Figure 3.**
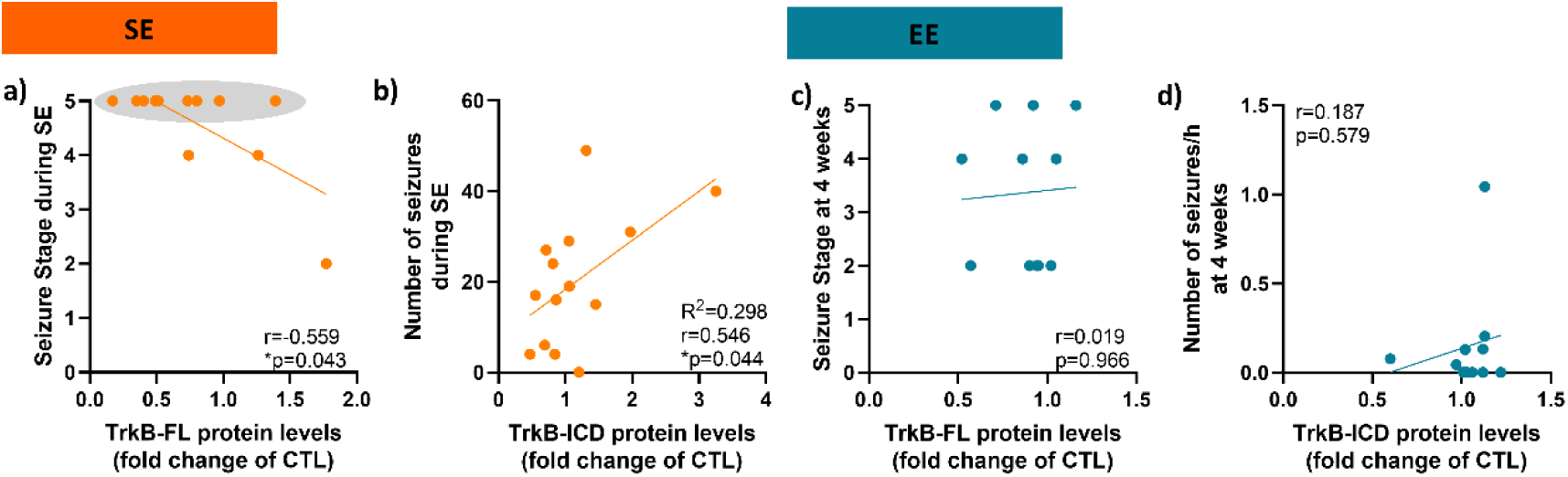
TrkB-FL levels are lower in hippocampal samples from animals treated with 10 mg/kg of KA and higher seizure stage during status epilepticus (SE). (a) Spearman correlation between the hippocampal TrkB-FL protein levels and seizure stage during SE (n=14). (b) Pearson correlation between the hippocampal TrkB-ICD protein levels and the number of seizures during SE (n=14). (c) Spearman correlation between the hippocampal TrkB-FL protein levels and seizure stage during established epilepsy (EE) (n=11). (d) Spearman correlation between the hippocampal TrkB-ICD protein levels and the number of seizures per hour during EE (n=11). Grey oval shape indicates all animals with stage 5 seizures.

In EE, no correlation was found between the seizure stage and TrkB-FL protein levels in the hippocampus of animals treated with 10 mg/kg of KA (Figure 3c). Also, no correlation was observed between TrkB-ICD levels and the number of seizures per hour (Figure 3d).

These findings highlight that in animals treated with 10 mg/kg of KA, TrkB-FL hippocampal levels decreased with the seizure stage during SE but not during EE.

### Cleavage of TrkB-FL occurs in KA-treated animals (10 mg/kg) with stage 5 seizures during SE

Taking into consideration that animals with a higher seizure stage during SE displayed lower levels of TrkB-FL, we further evaluated TrkB-FL cleavage in animals treated with 10 mg/kg of KA that had stage 5 seizures during SE (Figure 4). These animals had significantly lower TrkB-FL protein levels (p=0.004, Figure 4a) and higher TrkB-ICD/TrkB-FL ratio (p=0.035, Figure 4c) in the hippocampus compared with control animals. While no significant differences were found in TrkB-ICD protein levels in the hippocampus (Figure 4b) its levels were positively correlated with SE duration (p=0.004, Figure 4d) and with the number of seizures (p=0.010, Figure 4e) during SE. Regarding the cortex, no significant differences were found in the levels of TrkB-FL, TrkB-ICD, or TrkB-ICD/TrkB-FL ratio (Supplementary Figure 3a-c).

**Figure 4.**
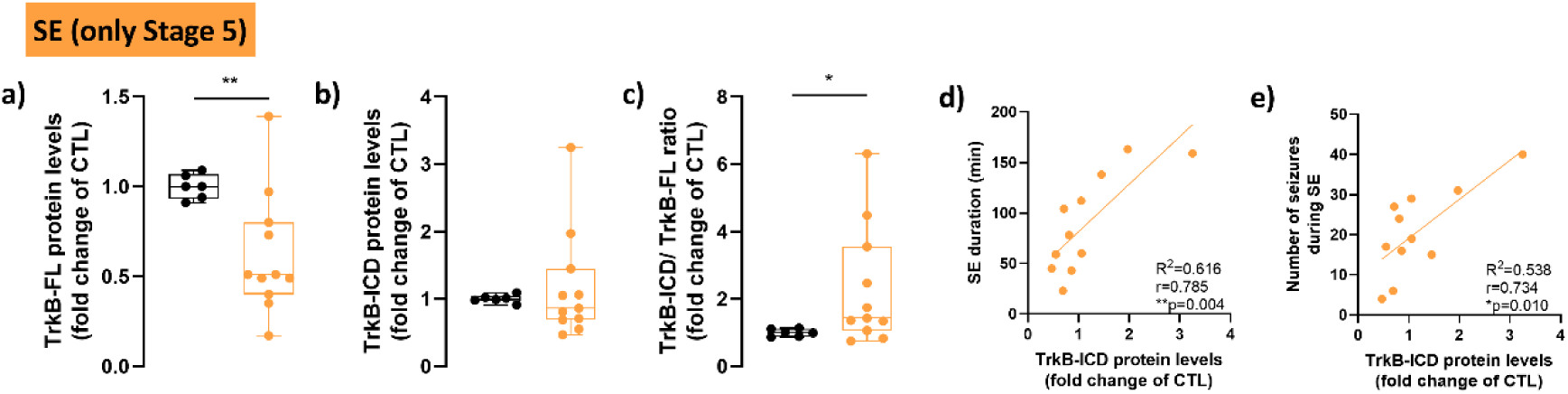
Cleavage of TrkB-FL during status epilepticus (SE) occurs in animals treated with 10 mg/kg of kainic acid (KA) displaying stage 5 seizures. (a) TrkB-FL levels, (b) TrkB-ICD levels, and (c) TrkB-ICD/TrkB-FL ratio from the hippocampus of control animals (black, n=6) and animals treated with 10 mg/kg of KA that experienced stage 5 seizures (orange, n=11) during SE. (d-e) Pearson correlation between the TrkB-ICD protein levels in the hippocampus and the SE duration (d) or the number of seizures during SE (e) (n=11). Statistical analysis was performed using an unpaired t-test with Welch’s correction (a-c). Results are shown as median, minimum, 25% and 75% percentile, and maximum. *p<0.05, **p<0.01, versus CTL.

These results emphasize that animals with higher TrkB-ICD levels experienced more seizures and longer SE, further suggesting the occurrence of TrkB-FL cleavage, particularly during SE with stage 5 seizures.

### KA-treated animals (10 mg/kg) present cleavage of TrkB-FL, neuronal death, mossy fiber sprouting, and memory impairment during EE

To understand whether TrkB-FL cleavage indeed persists throughout the process of epileptogenesis in animals treated with 10 mg/kg of KA, we performed a more detailed analysis of the previously described data (Figure 2d-f, Supplementary Figure 1d-f) by performing a direct comparison between control animals and animals treated with 10 mg/kg of KA instead of multiple comparisons with other doses. When comparing the hippocampus of animals treated with 10 mg/kg of KA with control animals, a significant increase in the TrkB-ICD protein levels was detected during EE (p=0.016, Figure 5b) along with an increase in the TrkB-ICD/TrkB-FL ratio (p=0.022, Figure 5c), although no differences were found in the levels of TrkB-FL (Figure 5a).

**Figure 5.**
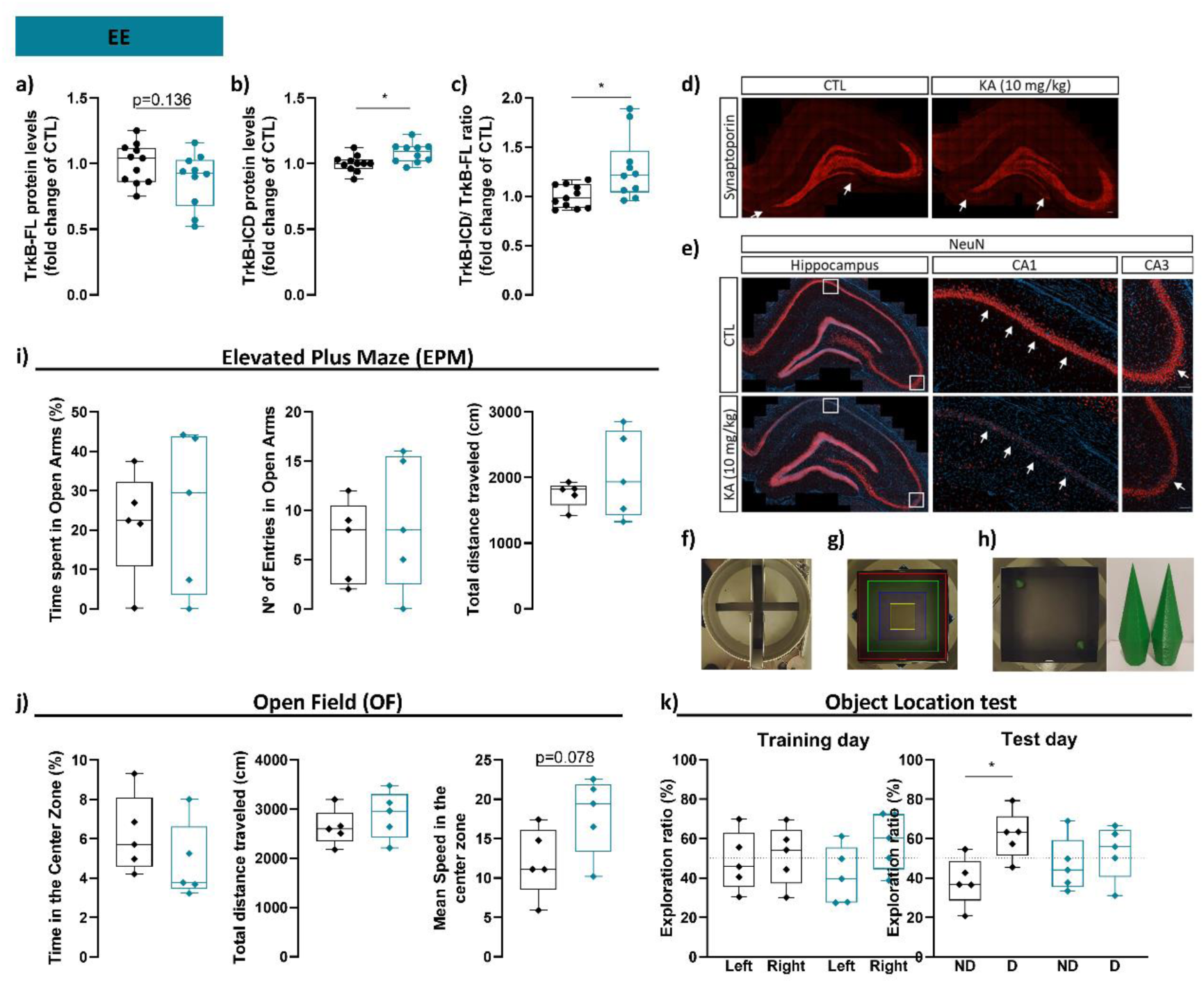
Animals treated with 10 mg/kg of kainic acid (KA) present TrkB-FL cleavage, mossy fiber sprouting, and neuronal death in the hippocampus, as well as, spatial memory impairment during established epilepsy (EE). (a) TrkB-FL levels, (b) TrkB-ICD levels, and (c) TrkB-ICD/TrkB-FL ratio from the hippocampus of control animals (black, n=11) and animals treated with 10 mg/kg of KA (blue, n=10) during EE. Immunohistochemical staining for synaptoporin (d) and NeuN (e) in the hippocampus of control animals and animals treated with 10 mg/kg of KA. White squares indicate the amplified CA1 and CA3 regions. Arrows indicate mossy fiber sprouting (d) or neuronal loss (e). Scale bar:100 µm. (f) Elevated plus maze. (g) Open field arena with virtual areas. (h) Objects used during the object location test and an example of its location during the test day. Evaluation of (i) elevated plus maze, (j) open field test, and (k) object location test in control animals (black, n=5) and animals treated with 10 mg/kg of KA (blue, n=5) during EE. Statistical analysis was performed using an unpaired t-test with Welch’s correction (a, c, i, and j), Mann-Whitney test (b), or paired t-test (k). Results are shown as median, minimum, 25% and 75% percentile, and maximum. *p<0.05 versus CTL. ND, No-displaced. D, displaced.

Simultaneously, in the cortex, a tendency to a decrease in TrkB-FL protein levels was depicted when compared with control animals during EE (p=0.069, Supplementary Figure 3d). Nevertheless, no differences were found in the cortical levels of TrkB-ICD or TrkB-ICD/TrkB-FL ratio in these animals (Supplementary Figure 3e and f).

To further identify the morphological alterations that occur during EE in the mTLE model used, mossy fiber sprouting, neuronal death, and memory were evaluated. Animals treated with 10 mg/kg of KA revealed mossy fiber sprouting (Figure 5d) together with a remarkable loss of NeuN^+^ mature neurons in the pyramidal layer of CA1 and CA3 hippocampal regions (Figure 5e). Thus, suggesting aberrant excitatory circuitry formation and neuronal death during EE.

During behavioral tests, the animals treated with 10 mg/kg of KA showed no anxiety-like behavior in the elevated plus maze (EPM) test, as indicated by similar time spent in the open arms, number of entries into the open arms, and total distance traveled compared with control animals (Figure 5i). Similarly, no differences were observed in locomotor activity in the open field (OF) test, as assessed by the time spent in the center zone and the total distance traveled in the arena (Figure 5j). Nevertheless, these animals displayed a tendency for increased mean speed in the center zone of the arena (p=0.078, Figure 5j).

Hippocampal-dependent long-term memory was assessed by the object location test. On the training day, animals from both conditions spent a similar amount of time exploring both objects (Figure 5k), indicating no alterations in the exploratory drive. On the test day, performed 24h after the training, control animals spent more time exploring the displaced object (p=0.049, Figure 5k). Contrastingly, animals treated with 10 mg/kg of KA did not spend more time exploring the displaced object, demonstrating a lack of preference for the object in the new location (Figure 5k), and therefore, a lack of memory for the object location observed 24h before, during training.

Altogether, these data suggest that the animals treated with 10 mg/kg of KA present aberrant excitatory circuitry, neuronal death, and impaired long-term spatial memory and discrimination during EE.

### Cleavage of TrkB-FL occurs in hippocampal samples from patients with refractory epilepsy

To further investigate the translational value of the obtained data, we analyzed human samples from autopsies of control individuals with no history of neurological diseases and from hippocampal resection of patients with refractory epilepsy (details in Supplementary Table 3 and Supplementary Table 4). Patients with epilepsy displayed significantly lower TrkB-FL protein levels compared with control individuals (p=0.0008, Figure 6a and b). Concomitantly, patients with epilepsy demonstrated significantly higher TrkB-ICD protein levels (p=0.0011, Figure 6a and c) and higher TrkB-ICD/TrkB-FL ratio (p=0.0001, Figure 6a and d) than control individuals, indicating that the cleavage of TrkB-FL also occurs in patients with refractory epilepsy.

**Figure 6.**
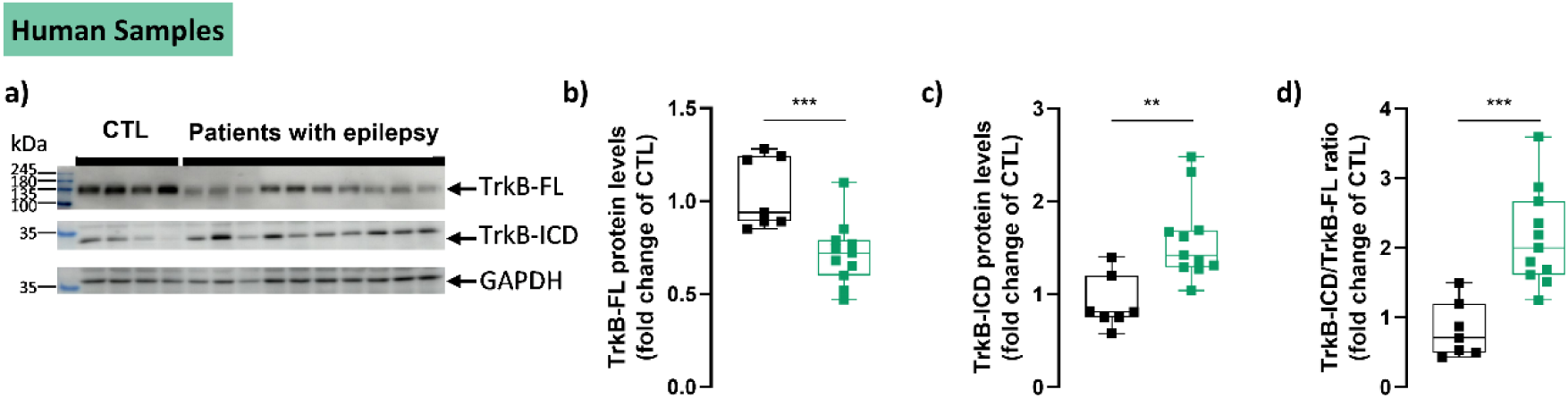
Cleavage of TrkB-FL occurs in human hippocampal samples from patients with refractory epilepsy. (a) Representative western blot bands of TrkB-FL (∼145 kDa), TrB-ICD (∼32 kDa), and GAPDH (∼37 kDa) of post-mortem tissue from control (CTL) humans and of resected epileptic tissue from patients with refractory epilepsy. (b) TrkB-FL protein levels, (c) TrkB-ICD protein levels, and (d) TrkB-ICD/TrkB-FL ratio of post-mortem tissue from control patients (black, n=7) and of epileptic tissue resected from patients with refractory epilepsy (green, n=11). Statistical analysis was performed using the Mann-Whitney test (b) or unpaired t-test with Welch’s correction (c and d). Results are shown as median, minimum, 25% and 75% percentile, and maximum. **p<0.01, ***p<0.001, versus CTL.

### TrkB-ICD overexpression induces long-term memory impairment in healthy rodents

Since TrkB-ICD has been shown to have a negative impact on synaptic plasticity (23), this fragment was overexpressed in the hippocampus of healthy rodents and its effects were assessed on memory, learning, and both short- and long-term synaptic plasticity. Lentiviruses overexpressing either TrkB-ICD or a control fluorescent protein (eGFP) (Figure 7a) were injected into the hippocampi of C57Bl/6J mice (Figure 8a) and Sprague-Dawley rats (Supplementary Figure 4a).

**Figure 7.**
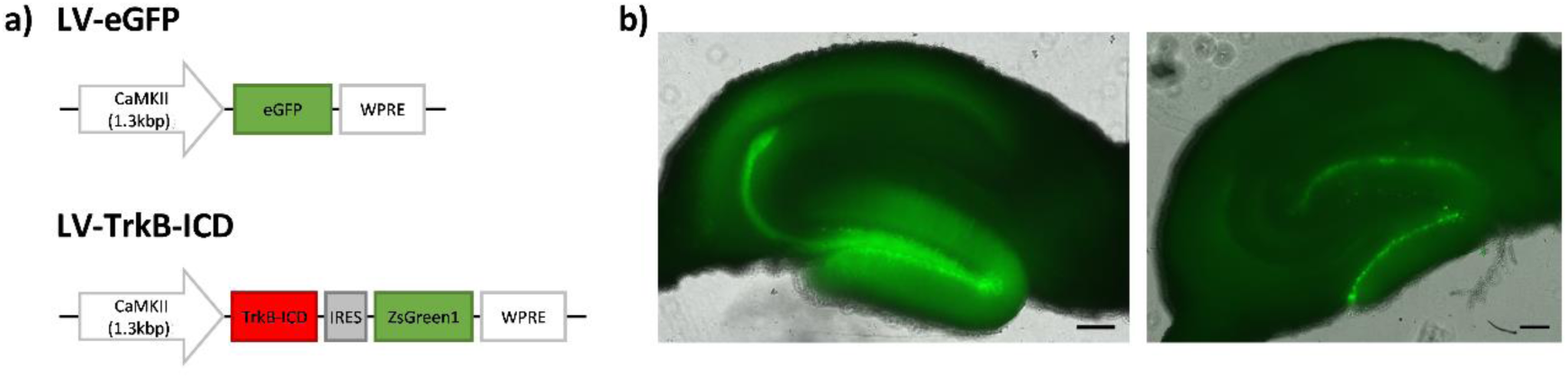
Design and overexpression of lentiviral vectors. (a) Constructs of lentiviral vectors expressing eGFP or TrkB-ICD. (b) Representative immunofluorescence images showing eGFP fluorescence from a slice of an animal injected with eGFP lentiviral vector (left panel) and ZsGreen fluorescence from a slice of an animal injected with TrkB-ICD lentiviral vector (right panel). Images were acquired with a widefield fluorescence microscope (Axiovert 200, ZEISS). Scale bar: 100 µm.

**Figure 8.**
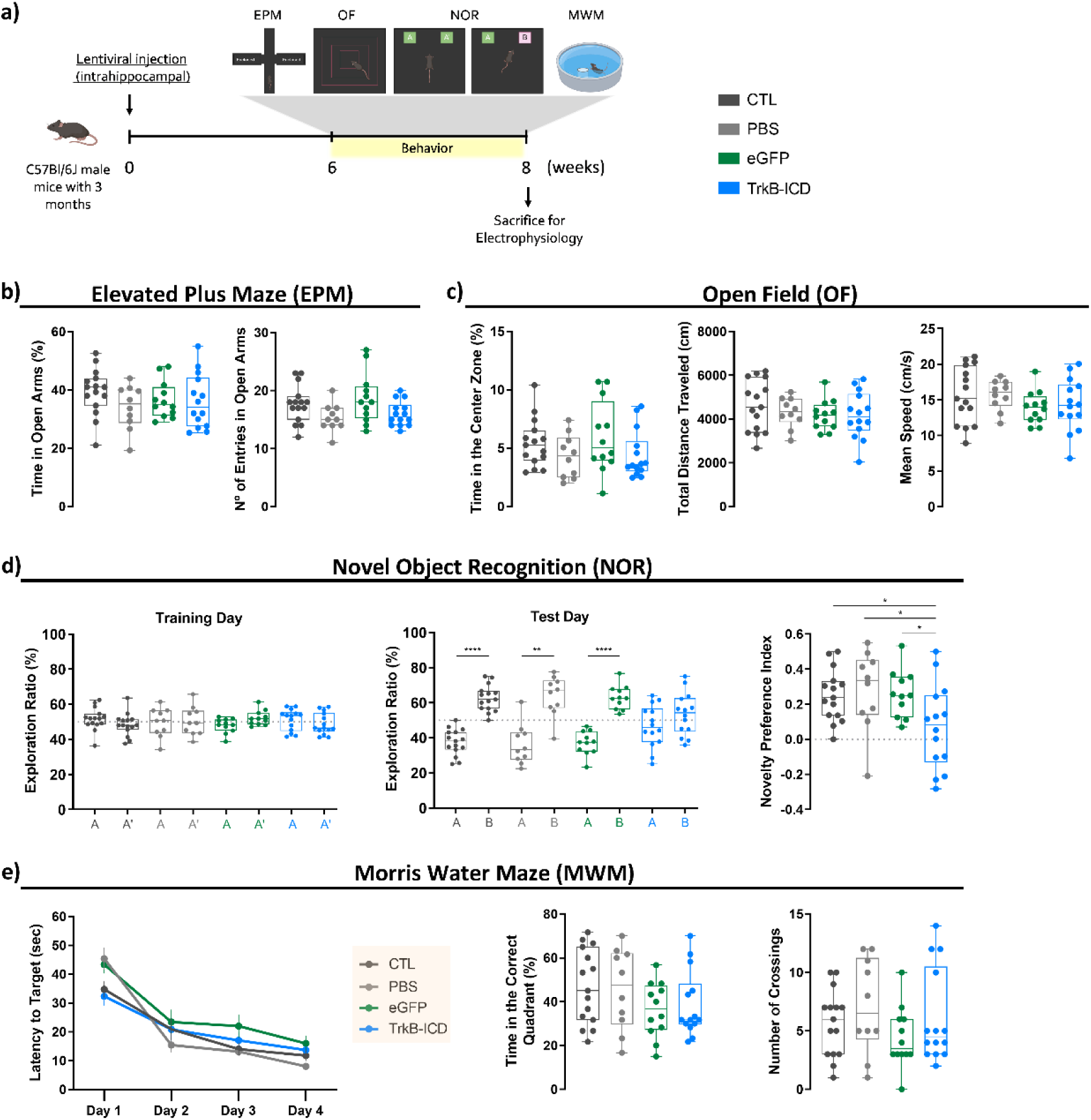
C57Bl/6J mice overexpressing TrkB-ICD exhibit long-term memory impairment. (a) Experimental design for animals injected with eGFP or TrkB-ICD lentiviral vectors, injected with a vehicle solution (phosphate-buffered saline, PBS), or not subjected to surgery. C57Bl/6J mice performed behavioral tests 6 weeks after lentiviral injection and were subsequently sacrificed for electrophysiological studies. (b-e) Evaluation of the different behavioral tests: (b) elevated plus maze, (c) open field, (d) novel object recognition, and (e) Morris water maze. The assessed groups included: control animals with no surgery (CTL, dark grey, n=15), animals injected with PBS (grey, n=10), animals injected with lentivirus for eGFP overexpression (green, n=12, except for NOR: n=11), and animals injected with lentivirus for TrkB-ICD overexpression (blue, n=14). Statistical analysis was performed using one-way ANOVA followed by Dunnett’s multiple comparisons tests (b, c, novelty preference index in (d), and time in the correct quadrant and the number of crossings in (e)), paired t-test (exploration ratio in (d)) and mixed-effects analysis (latency to target in (e)). Results are shown as median, minimum, 25% and 75% percentile, and maximum. *p<0.05, **p<0.01, ***p<0.001, ****p<0.0001.

To confirm lentiviral expression in injected mice, fluorescent proteins (eGFP and ZsGreen) were observed prior to electrophysiological recordings (Figure 7b). Rats injected with the TrkB-ICD lentiviral vector showed ZsGreen fluorescence and pronounced staining with Pan-Trk antibody, mainly in the innermost layer of the granule cell layer 1 (GCL 1) within the dentate gyrus (Supplementary Figure 5a). In contrast, animals injected with the eGFP lentiviral vector exhibited eGFP fluorescence but did not show enhanced staining for Pan-Trk (Supplementary Figure 5b). Since the Pan-Trk antibody binds to the intracellular domain of TrkB-FL, thus recognizing both TrkB-ICD and TrkB-FL, we further demonstrated, using an antibody specific to the extracellular domain of TrkB-FL, that Pan-Trk predominantly labeled TrkB-ICD (Supplementary Figure 5c). These results indicate that TrkB-ICD was effectively overexpressed in these animals.

After 6 weeks of lentiviral overexpression, a battery of behavioral tests was conducted on animals overexpressing either eGFP or TrkB-ICD, as well as, on animals injected with the vehicle solution (PBS) or with no surgery (CTL).

During behavioral tests, mice did not exhibit anxiety-like behavior, as indicated by a similar number of entries and time spent in the open arms of the EPM (Figure 8b). Additionally, no differences were observed in locomotor activity, assessed by the time in the center zone, the total distance traveled, and the mean speed during OF (Figure 8c). These findings suggest that TrkB-ICD overexpression does not affect anxiety-like behavior or locomotion.

Long-term recognition memory was assessed using the NOR test. On the training day, all groups — including the control mice (CTL, PBS, and eGFP) and the TrkB-ICD-overexpressing mice — spent similar amounts of time exploring both objects (A and A’, Figure 8c). On the test day, the control groups spent more time exploring the novel object (B), successfully discriminating it (CTL_AvsB_: p<0.0001, PBS_AvsB_: p=0.004, eGFP_AvsB_: p<0.0001, Figure 8d). However, TrkB-ICD-overexpressing mice failed to significantly discriminate the novel object, spending similar amounts of time exploring both objects (A and B, Figure 8d). The analysis of the NPI further supported this conclusion (TrkB-ICD_vs_CTL: p=0.039, TrkB-ICD_vs_PBS: p=0.032, TrkB-ICD_vs_eGFP: p=0.048, Figure 8d).

Thus, these results suggest that overexpression of TrkB-ICD in the hippocampus of healthy mice impairs long-term recognition memory.

Finally, hippocampal-dependent spatial memory was evaluated using the MWM test. During the 4-day acquisition phase, the latency to escape, measured by the time taken to reach the hidden platform, was similar across all groups (Figure 8e). In the probe test, the number of platform crossings and the time spent in the target quadrant were also similar across groups (Figure 8e). These results suggest no significant differences in long-term spatial memory among the experimental groups during this test.

To further validate these findings, a battery of behavioral tests was also conducted in Sprague-Dawley rats following lentiviral injection (Supplementary Figure 4). The OF test revealed no changes in locomotion between TrkB-ICD-expressing and eGFP-expressing rats (Supplementary Figure 4c). Surprisingly, TrkB-ICD-overexpressing rats exhibited a significant reduction in the number of entries into the open arms of the EPM (p=0.021, Supplementary Figure 4b), although no significant differences were observed in the time spent in the open arms of the EPM or in the central zone of the OF arena (Supplementary Figure 4b), suggesting a more anxiety-like behavior.

Subsequently, we performed the object location test to assess hippocampus-dependent long-term memory specifically. On the training day of the object location test, eGFP-overexpressing rats spent an equal amount of time exploring both objects (Supplementary Figure 4d). In contrast, TrkB-ICD-overexpressing rats spent more time exploring the object on the right (p=0.040, Supplementary Figure 4d). However, there was no significant preference for turning tendency or for initially choosing the object on the right side compared to the eGFP-expressing rats (Supplementary Figure 4d). Remarkably, as observed in mice during the NOR test, TrkB-ICD-expressing rats failed to discriminate the displaced object on the test day, spending similar amounts of time exploring both objects, while eGFP-expressing rats spent more time exploring the displaced object (p=0.038, Supplementary Figure 4d). These results suggest that TrkB-ICD overexpression impairs hippocampus-dependent long-term memory.

### TrkB-ICD overexpression induces dysfunctional hippocampal long-term potentiation in healthy mice

Following the behavioral assessment in mice, we analyzed short-term plasticity (by measuring PTP) and long-term plasticity (by measuring LTP). Short- and long-term synaptic plasticity are fundamental mechanisms underlying learning and memory. To investigate these mechanisms, fluorescently labeled dorsal hippocampal slices from mice expressing either eGFP or TrkB-ICD, along with all slices from CTL and PBS-injected mice, were used for ex vivo electrophysiological recordings.

The PTP magnitude showed that neither PBS injection nor eGFP expression significantly affected basal PTP levels compared with CTL mice (Figure 9b). Moreover, TrkB-ICD overexpression did not significantly alter this form of short-term plasticity (Figure 9b).

**Figure 9.**
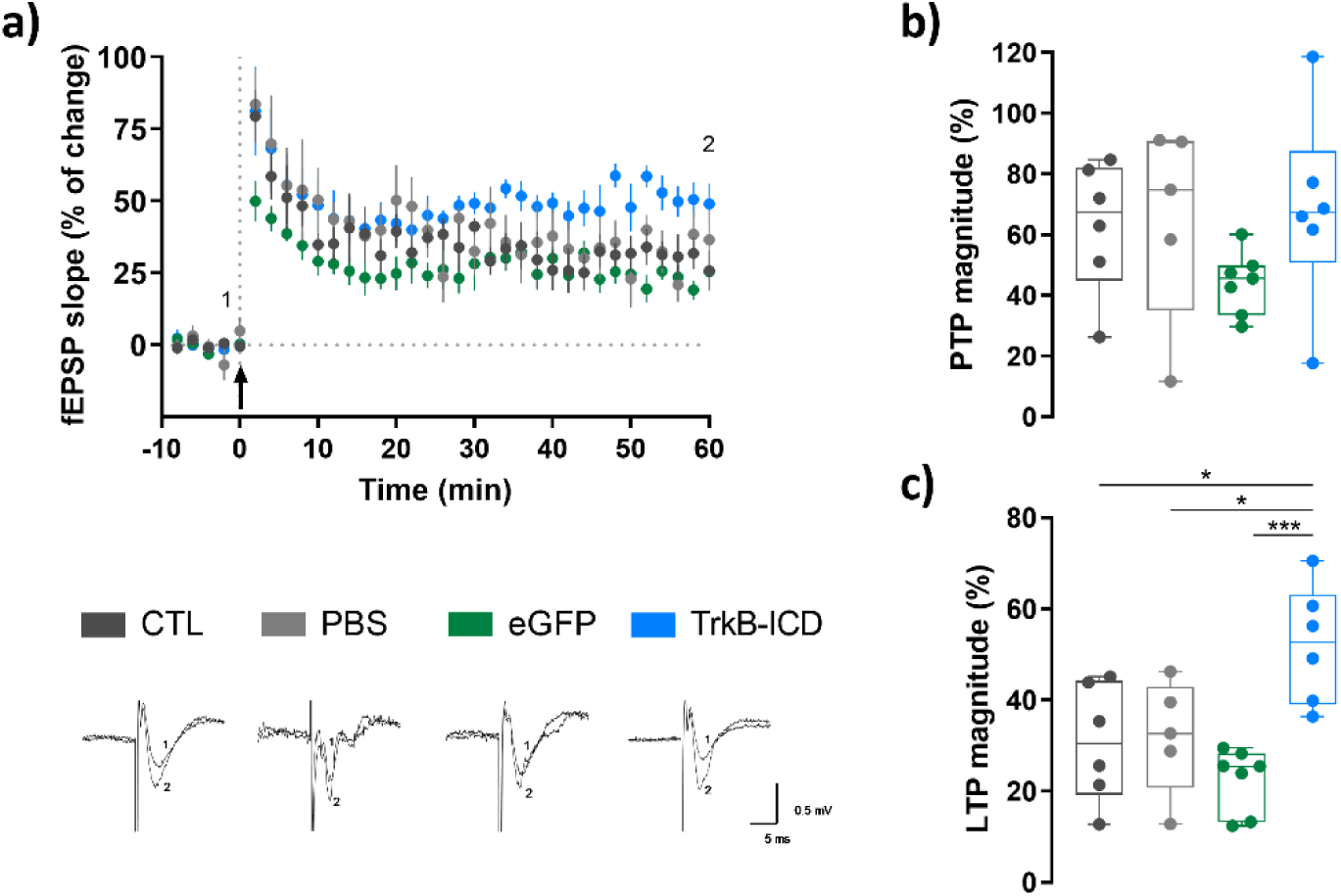
TrkB-ICD overexpression in the hippocampus of C57Bl/6J mice promoted a dysfunctional increase in long-term synaptic plasticity, without affecting short-term synaptic plasticity. (a) Time courses of averaged normalized changes in field excitatory postsynaptic potentials (fEPSP) slope after delivery (0 min) of a weak θ-burst (3×3) in all conditions (CTL, dark grey, n=6; PBS, grey, n=5; eGFP, green, n=7; TrkB-ICD, blue n=6). Representative traces were recorded before (1) and after (2) long-term potentiation (LTP) induction from the experimental group identified above each trace. (b) Post-tetanic potentiation (PTP) magnitude after θ-burst stimulation (change in fEPSP slope at 0-6 minutes (recorded in a), compared to baseline) in all conditions. (c) LTP magnitudes from the recordings in (a) (change in fEPSP slope at 50-60 min compared to baseline). Statistical analysis was performed using a one-way ANOVA followed by Dunnett’s multiple comparisons test (b and c). Results are shown as median, minimum, 25% and 75% percentile, and maximum. *p<0.05, ***p<0.001, versus TrkB-ICD.

Regarding long-term plasticity, neither PBS injection nor eGFP expression affected LTP magnitude compared with CTL mice (Figure 9c). Remarkably, TrkB-ICD overexpression significantly increased the LTP magnitude compared to the control groups (TrkB-ICD_vs_CTL: p=0.011, TrkB-ICD_vs_PBS: p=0.023, TrkB-ICD_vs_eGFP: p=0.0005, Figure 9c).

In summary, these findings suggest that TrkB-ICD overexpression markedly exacerbates long-term synaptic plasticity without affecting short-term plasticity.

## Discussion

In this study, we demonstrated the cleavage of TrkB-FL and the subsequent formation of TrkB-ICD both in an in vivo rat model of SE and epilepsy and in samples obtained from patients with refractory epilepsy. During SE induced by KA, TrkB-FL cleavage mainly occurs in animals that reach the highest seizure stage on the modified Racine scale (stage 5), with TrkB-ICD formation being proportional to the number of seizures. Furthermore, TrkB-FL cleavage is also observed in rats with EE and is prominently detected in the hippocampus of patients with refractory epilepsy. Furthermore, our findings indicate that the intrahippocampal overexpression of TrkB-ICD leads to long-term memory impairment. Therefore, we suggest a mechanistic role for TrkB-ICD, highlighting its detrimental impact on long-term memory.

KA, a drug widely used in epilepsy research (32,33), was used to induce the experimental model (34,35). Considering the variability of published methods in the literature (5,34,35), different dose regimens were tested to identify the most adequate approach to induce the mTLE model. We found that the dose of 10 mg/kg of KA is the most effective, as it consistently induced stage 5 seizures during SE, and SRCS in EE. The observed variability in seizure severity was not regarded as a limitation but rather as an opportunity to further investigate the mechanism in focus. Additionally, this methodology allowed us to work with a larger number of animals without introducing confounding factors such as the use of anesthetics (36), which are required when KA is directly administered in the brain (32,37).

To validate the model, we assessed whether the animals treated with KA exhibited key characteristics of epilepsy during EE. Specifically, we observed neuronal death, consistent with findings reported by others during EE after KA (i.p.) (38,39), pilocarpine (40), or kindling (41). Mossy fiber sprouting was also evident, aligning with other works demonstrating this phenomenon during EE after KA (i.p.) (39), pilocarpine (40), or kindling (42). Additionally, animals treated with KA exhibited long-term memory impairment, as described in studies using KA (43) or pilocarpine (44).

The role of the BDNF/TrkB-FL signaling in epilepsy remains puzzling as it may have pro-epileptic effects through the PLCγ1-mediated pathway (45), or anti-epileptic actions (13,14). In an experimental model using an i.p. administration of KA (12 mg/kg), mRNA and protein levels of BDNF were increased, peaking 24h later (5). Moreover, the authors also showed that BDNF protein levels were higher in animals that experienced seizures compared with non-seizing animals (5). This finding was also seen in other models of epilepsy such as in pilocarpine (9,11) or kindling (46) models. Whether the increase in mRNA and protein levels of BDNF detected in those works constitute a feedback response to decreased BDNF signaling due to cleavage of TrkB-FL receptors was not addressed and requires further clarification.

It was previously shown that TrkB-FL cleavage contributes to neuronal death (21) and memory impairment (23), in excitotoxic contexts. This cleavage not only implies the decrease of TrkB-FL levels but also the formation of two fragments: a new truncated isoform and an intracellular domain (TrkB-ICD) (18). Indeed, increased levels of truncated TrkB isoforms were found after KA (5,6,8,21) or pilocarpine (10) administration or after kindling (12). However, to date, there is no data concerning TrkB-ICD formation in the context of epilepsy or focusing on the relationship between the seizure severity and the levels of TrkB-FL or TrkB-ICD. Importantly, due to the pathophysiological implications, a comparison between TrkB-FL cleavage in SE and EE is of relevance but was missing.

Our results show that the degree of TrkB-FL cleavage depends on the administered dose of KA and the seizure stage reached. Accordingly, during SE, only the animals treated with 10 mg/kg of KA and that reached stage 5, on the modified Racine scale, showed a significant decrease in TrkB-FL protein levels. The decrease of TrkB-FL protein levels was described by others, hours to days after KA (7,8) or pilocarpine (9) administration, as well as in in vitro models of SE induced by KA or Mg^2+^-free medium (21,47). Contrastingly, TrkB-FL mRNA levels were described to be maintained until 7 days after KA (5,6), or increased hours after pilocarpine or kindling (11,12). Contrary to our findings in EE, which revealed no significant changes in TrkB-FL protein levels, another study reported an increase in TrkB-FL protein levels in EE induced by pilocarpine (10). Intriguingly, the same authors also demonstrated a decrease in BDNF protein levels more than 60 days after pilocarpine administration (10).

In this work, using an antibody that recognizes the intracellular domain of TrkB-FL, as in our previous studies (18,22), we revealed for the first time the formation of TrkB-ICD in an in vivo model of SE and epilepsy. It is noteworthy that TrkB-ICD protein levels were also detected in control animals, suggesting that a basal TrkB-FL cleavage may occur under physiological conditions. Importantly, we have shown an increase in TrkB-ICD/TrkB-FL ratio in animals during SE and that TrkB-ICD formation is associated with an increase in seizure severity during this period. Moreover, we demonstrated an increase in TrkB-ICD protein levels and in the TrkB-ICD/TrkB-FL ratio in animals with EE, suggesting the cleavage of the TrkB-FL throughout the epileptogenesis process.

One could argue that the cleavage of TrkB-FL might be considered an artifact of the model, as KA, functioning as a glutamate analog, could potentially increase the calcium influx (48,49), thereby triggering the cleavage of TrkB-FL, as previously observed in vitro (18). However, if this were the case, one would expect similar degrees of cleavage in both the cortex and hippocampus, which contradicts our data showing more consistent cleavage in the hippocampus than in the cortex. Additionally, the decreased levels of TrkB-FL and the increased levels of TrkB-ICD detected in hippocampal samples from patients with refractory epilepsy further support that these findings are not artifacts. TrkB-FL cleavage was consistently detected in all samples of these patients with the most severe cases of epilepsy. However, it remains unclear whether this mechanism occurs in less severe cases, in seizure-free cases managed with antiseizure drugs, or in other epilepsy syndromes in humans. This uncertainty arises from the inability to collect samples in such cases or the challenges associated with obtaining them, particularly in rare epilepsy syndromes. Our findings in human samples contrast with a previous study (3) reporting increased TrkB-FL protein and mRNA levels in patients with intractable epilepsy compared with hippocampal samples collected from autopsies.

Given the significant influence of age on TrkB-FL levels - where older animals generally exhibit lower levels compared to younger counterparts under healthy conditions (50) - it is plausible that age may contribute to these observed differences. In our study, control samples were obtained from individuals with a median age twice that of the epilepsy patients. Despite this, control samples displayed higher TrkB-FL protein levels than those from patients with epilepsy, suggesting that the observed differences are more likely associated with epilepsy itself. Notably, the age of the patients reported by Hou and colleagues (3) is similar between conditions and similar to the age of the patients with epilepsy in our study. Thus, age does not appear to account for the differences in TrkB-FL levels observed between studies. Another aspect that has to be taken into account and may impact the results is antiseizure medication. Besides reporting an increase in TrkB-FL levels in patients with refractory epilepsy, Hou and colleagues also reported the impact of valproate in decreasing these levels (3). Therefore, the differences among the studies might be due to differences in the pharmacological treatments of the patients.

Here, our findings also provide the first evidence of the detrimental impact of TrkB-ICD overexpression on the long-term memory of healthy rodents. Using lentiviral vectors, we demonstrated that TrkB-ICD overexpression leads to long-term memory impairment in the NOR and object location tests and induces a dysfunctional increase in LTP, a key molecular basis of learning and memory. These findings build upon our recently published study, which examined the effects of TrkB-ICD overexpression in vitro, using the same lentiviral vectors (23). In this recently published study, overexpression of TrkB-ICD in primary neuronal cultures resulted in several changes: (i) a significant decrease in the number of dendritic spines without affecting cell survival, (ii) a hyperpolarized resting membrane potential and an increase in the frequency of miniature excitatory postsynaptic currents (mEPSCs), (iii) transcriptome alterations in genes encoding for synaptic proteins (23). Curiously, one of the reported genes altered by TrkB-ICD is *Npy2r* (23), which encodes the neuropeptide Y receptor Y2. Increased mRNA levels of this receptor are well-documented in the literature for both SE and established epilepsy in rodent models and human patients (51–54). Thus, we hypothesize that TrkB-ICD-induced Y2 receptor upregulation works as a neuroprotective mechanism to counteract the effects of TrkB-ICD, as the Y2 receptor appears to inhibit glutamate release (51).

In behavioral testing of animals overexpressing TrkB-ICD, neither mice nor rats showed locomotor alterations. Although rats exhibited a significant reduction in open-arm entries in the EPM, pronounced anxiety-like behavior was not observed, consistent with findings in mice. In novel object recognition tests in mice, TrkB-ICD overexpression led to impaired long-term memory, and subsequent object location tests in rats confirmed that this impairment was hippocampus-dependent. However, the MWM test in mice did not reveal any alteration in the hippocampal-dependent memory. One explanation could be the stress associated with the MWM test, which might mask memory impairments compared to less stressful models (55). While long-term memory deficits were evident in novel object recognition and object location tests, these effects do not appear to result from decreased long-term learning and memory mechanisms. Instead, our results suggest that TrkB-ICD may lead to a dysfunctional increase in synaptic plasticity.

The results described here reveal a new mechanism in epileptogenesis that may elucidate the dual role of BDNF/TrkB-FL signaling in epilepsy. This mechanism, also identified in patients with AD (18), may explain their increased susceptibility to seizures (56). In addition, neuronal hyperexcitability has been shown to accelerate AD progression (57). Moreover, the formation of TrkB-ICD could be linked to cognitive impairments observed in patients with epilepsy and AD (58–60). To conclude, this newly identified mechanism in epilepsy can be responsible for drug resistance and represents a promising therapeutic target for the development of next-generation antiseizure drugs. Strategies such as using a drug to prevent the cleavage of TrkB-FL or to inhibit the detrimental effects of TrkB-ICD on memory, or potentially a combination of both, could offer novel approaches to controlling seizures in cases of refractory epilepsy or serve as disease-modifying strategies, preventing epilepsy progression.

## Conclusion

In this work, we provide evidence for the cleavage of TrkB-FL and the formation of TrkB-ICD in the mTLE animal model and human patients with epilepsy. We demonstrate that this mechanism persists throughout epileptogenesis and is dependent on seizure severity. Additionally, we clarify the role of TrkB-ICD in long-term memory.

The discovery of TrkB-ICD formation as a result of TrkB-FL cleavage in epilepsy has provided new insights into the disease’s underlying mechanisms and revealed a potential new pharmacological target for epilepsy treatment.

## Acknowledgments

This work was supported by *Santa Casa da Misericórdia de Lisboa* [grant numbers MB37-2017, MB-35-2021]; International Brain Research Organization (IBRO) Early Career Award, European Union’s Horizon 2020 research and innovation program H2020-WIDESPREAD-05-2020-Twinning (EpiEpinet) [grant number: 952455], and *Bolsas CHLN/FMUL* under the “*Educação pela Ciência*” – GAPIC Program [grant number: 20210019 and 20220006]. *Fundação para a Ciência e Tecnologia* (FCT) supported L.R-R. (PD/BD/150344/2019), J.F-G. (SFRH/PD/BD/114441/2016), S.L.P (PD/BD/150341/2019), F.F.R. (IMM/CT/35-2018), C.M-L. (PD/BD/118238/2016), F.M.M. (IMM/CT/8-2018), R.F.B. (PD/BD/114337/2016), C.B.F. (PD/BD/128390/2017), S.R.T. (PD/BD/128091/2016), M.F-M (PD/BD/10313/2022) and V.H.P. (2021.01812.CEECIND/ CP1656/CT0014).

The authors are grateful to all people who contributed to this work in particular from the Biobank (Ângela Afonso), Rodent (Iolanda Moreira, Daniel Gomes da Costa, Patricia Almeida, Márcia Silva), Bioimaging (José Rino, António Temudo, Ana Nascimento), and Histology and Comparative Pathology (Ana Rita Pires, Ana Margarida Biscaia and Ana Margarida Cristovão), Facilities at Instituto de Medicina Molecular João Lobo Antunes (iMM). The authors of this paper also want to acknowledge Joana Coelho for the suggestions regarding the model of epilepsy and behavioral tests and Diana Sousa Correia for helping with animal monitoring.

Graphical Abstract was created in BioRender. Sebastião, A. (2024) https://BioRender.com/e62o244.

## Authors contribution

LR-R: Conceptualization (supporting); Data Curation; Formal analyses; Funding Acquisition (supporting); Investigation (lead, all experiments except mice experiments); Project Administration; Visualization; Writing – Original Draft Preparation (lead). SL-P: Funding Acquisition (supporting); Investigation (supporting, perfusions and behavioral tests in rats); Writing – Review & Editing (supporting). JF-G: Funding Acquisition (supporting); Investigation (lead, on mice experiments); Writing – Review & Editing (supporting). RV: Investigation (supporting, lentiviral production); Writing – Review & Editing (supporting). FRR: Funding Acquisition (supporting); Investigation (supporting, mice surgeries); Writing – Review & Editing (supporting). CM-L: Funding Acquisition (supporting); Investigation (supporting, mice surgeries and electrophysiology); Writing – Review & Editing (supporting). FMM: Funding Acquisition (supporting); Investigation (supporting, behavioral tests in mice). RFB: Funding Acquisition (supporting); Investigation (supporting, electrophysiology). CBF: Funding Acquisition (supporting); Investigation (supporting, electrophysiology). SRT: Funding Acquisition (supporting); Investigation (supporting, electrophysiology). MFM: Funding Acquisition (supporting); Investigation (supporting, mice experiments). JM: Investigation (supporting, lentiviral production). EC: Supervision (supporting, lentiviral production). VHP: Formal Analyses (supporting); Writing – Review & Editing (supporting). AMS: Funding Acquisition (supporting); Writing – Review & Editing (equal). EA: Resources (lead, post-mortem samples); Writing – Review & Editing (supporting). ARC: Resources (lead, samples from patients with epilepsy). CB: Resources (supporting, samples from patients with epilepsy); Writing – Review & Editing (supporting). SX: Conceptualization (lead); Funding Acquisition (lead); Supervision (equal); Writing – Review & Editing (lead). MJD: Conceptualization (lead); Funding Acquisition (lead); Supervision (equal); Writing – Review & Editing (lead). All authors read, revised, and approved the final manuscript.

## Data availability

Data will be made available on request.

## Supplementary Tables

**Supplementary Table 1.** Methodology used in Sprague-Dawley male rats injected with saline or kainic acid sacrificed during status epilepticus.

| Animal | Condition | Time of sacrifice | SE Stage | Nº of seizures during SE | WB in the right hippocampus | WB in the right cortex |
| --- | --- | --- | --- | --- | --- | --- |
| S1 | Saline | 3h | - | - | + | + |
| S2 | Saline | 3h | - | - | + | + |
| S5 | Saline, 3 i.p. | 3h | - | - | + | + |
| S6 | Saline, 3 i.p. | 3h | - | - | + | + |
| S10 | Saline | 3h | - | - | + | + |
| S11 | Saline | 3h | - | - | + | + |
| S14 | Saline | 3h | - | - | + | + |
| S15 | Saline | 3h | - | - | + | + |
| K7 | 5 mg/kg, 3 i.p. | 2h51 | 2 | 0 | + | + |
| K8 | 5 mg/kg, 3 i.p. | 3h02 | 2 | 0 | + | + |
| K9 | 5 mg/kg, 3 i.p. | 3h26 | 1 | 0 | + | + |
| K10 | 5 mg/kg, 3 i.p. | 6h33 | 5 | 27 | + | + |
| K13 | 5 mg/kg, 1 i.p. | 3h53 | 2 | 0 | + | + |
| K14 | 5 mg/kg, 1 i.p. | 2h40 | 5 | 32 | + | + |
| K15 | 5 mg/kg, 1 i.p. | 4h08 | 1 | 0 | + | + |
| K19 | 8.5 mg/kg | 3h27 | 2 | 0 | + | + |
| K20 | 8.5 mg/kg | 3h59 | 4 | 2 | + | + |
| K21 | 8.5 mg/kg | 2h47 | 5 | 35 | + | + |
| K22 | 8.5 mg/kg | 2h03 | 5 | 10 | + | + |
| K26 | 8.5 mg/kg | 1h32 | 5 | 13 | + | + |
| K1 | 10 mg/kg | 2h33 | 5 | 16 | + | + |
| K2 | 10 mg/kg | 2h23 | 5 | 19 | + | + |
| K3 | 10 mg/kg | 2h14 | 5 | 4 | + | + |
| K4 | 10 mg/kg | 4h09 | 5 | 15 | + | + |
| K27 | 10 mg/kg | 3h11 | 5 | 27 | + | + |
| K28 | 10 mg/kg | 3h42 | 5 | 40 | + | + |
| K29 | 10 mg/kg | 4h00 | 5 | 31 | + | + |
| K30 | 10 mg/kg | 2h19 | 5 | 17 | + | + |
| K31 | 10 mg/kg | 1h38 | 4 | 4 | + | + |
| K33 | 10 mg/kg | 1h49 | 5 | 6 | + | + |
| K34 | 10 mg/kg | 2h50 | 5 | 24 | + | + |
| K35 | 10 mg/kg | 3h04 | 5 | 29 | + | + |
| K36 | 10 mg/kg | 3h35 | 2 | 0 | + | + |
| K37 | 10 mg/kg | 3h52 | 4 | 49 | + | + |
| K42 | 10 mg/kg | 3h | 5 | - | - | - |
| K48 | 10 mg/kg | 2h17 | 5 | - | - | - |
| K49 | 10 mg/kg | 3h | 5 | - | - | - |

Abbreviations: S, Saline. K, Kainic acid. SE, status epilepticus. i.p., intraperitoneal injection. +, assessed. -, not assessed.

**Supplementary Table 2.** Methodology used in Sprague-Dawley male rats injected with saline or kainic acid sacrificed during established epilepsy (EE).

| Animal | Condition | Time of sacrifice | SE Stage | Nº of seizures during SE | EE Stage | Nº of seizures during EE | WB in the right hippo. | WB in the right cortex | IHC in the left hippo. | Behavior |
| --- | --- | --- | --- | --- | --- | --- | --- | --- | --- | --- |
| S3 | Saline | 4 w | - | - | - | - | + | + | - | - |
| S4 | Saline | 4 w | - | - | - | - | + | + | - | - |
| S7 | Saline, 3 i.p. | 4 w | - | - | - | - | + | + | - | - |
| S8 | Saline, 3 i.p. | 4 w | - | - | - | - | + | + | - | - |
| S9 | Saline, 3 i.p. | 4 w | - | - | - | - | + | + | - | - |
| S12 | Saline | 4 w | - | - | - | - | + | + | - | - |
| S13 | Saline | 4 w | - | - | - | - | + | + | - | - |
| S16 | Saline | 4 w | - | - | - | - | + | + | - | - |
| S17 | Saline | 4 w | - | - | - | - | + | + | - | - |
| S18 | Saline | 5 w | - | - | - | - | + | + | - | + |
| S19 | Saline | 5 w | - | - | - | - | + | + | - | + |
| S20 | Saline | 5 w | - | - | - | - | + | - | - | + |
| S24 | Saline | 5 w | - | - | - | - | + | - | - | + |
| S25 | Saline | 5 w | - | - | - | - | + | + | + | + |
| K11 | 5 mg/kg, 3 i.p. | 4 w | 2 | 0 | 4 | - | + | + | - | - |
| K12 | 5 mg/kg, 3 i.p. | 4 w | 2 | 0 | 4 | - | + | + | - | - |
| K16 | 5 mg/kg, 1 i.p. | 4 w | 2 | 0 | 2 | 0 | + | + | - | - |
| K17 | 5 mg/kg, 1 i.p. | 4 w | 2 | 0 | 2 | 0 | + | + | - | - |
| K18 | 5 mg/kg, 1 i.p. | 4 w | 2 | 0 | 2 | 0 | + | + | - | - |
| K23 | 8.5 mg/kg | 4 w | 2 | 0 | 2 | 0 | + | + | - | - |
| K24 | 8.5 mg/kg | 4 w | 2 | 0 | 2 | 0 | + | + | - | - |
| K25 | 8.5 mg/kg | 4 w | 2 | 0 | 2 | 0 | + | + | - | - |
| K5 | 10 mg/kg | 4 w | 2 | 0 | 2 | 0 | + | + | - | - |
| K6 | 10 mg/kg | 4 w | 2 | 0 | 4 | 0.129 | + | + | - | - |
| K32 | 10 mg/kg | 4 w | 1 | 0 | 4 | 0.077 | + | + | - | - |
| K38 | 10 mg/kg | 4 w | 2 | 0 | 4 | 0.204 | + | + | - | - |
| K39 | 10 mg/kg | 1 d | 5 | - | - | - | - | - | - | - |
| K40 | 10 mg/kg | 4 w | 4 | 30 | 5 | 1.043 | + | + | - | - |
| K41 | 10 mg/kg | 4 w | 1 | 0 | 2 | 0 | + | + | - | - |
| K43 | 10 mg/kg | 5 w | 4 | 35 | 2 | 0 | + | + | - | + |
| K44 | 10 mg/kg | 5 w | 1 | 0 | 2 | 0 | + | + | - | + |
| K45 | 10 mg/kg | 5 w | 2 | 0 | 2 | 0 | + | + | - | + |
| K46 | 10 mg/kg | 5 w | 2 | 0 | 5 | 0.132 | + | + | - | + |
| K47 | 10 mg/kg | 5 w | 5 | 45 | 5 | 0.044 | + | + | + | + |

Abbreviations: S, Saline. K, Kainic acid. SE, status epilepticus. EE, established epilepsy. Hippo., hippocampus. i.p., intraperitoneal injection. d, day. w, weeks. +, assessed. -, not assessed.

**Supplementary Table 3.** Clinical information of patients with epilepsy who underwent hippocampal resection. Patients 1, 4, 6, and 7 had dual pathology with both hippocampal sclerosis and adjacent temporal lobe dysplasia (patients 1, 4 and 7) or mesial sclerosis and a temporal lobe DNT (patient 6).

| **SAMPLE** | **AGE** | **SEX** | **LESIONS** | **LATERALITY**<br>**OF**<br>**RESECTION** |
| --- | --- | --- | --- | --- |
| **EHS 1** | 6 | M | Hippocampal sclerosis/ FCD | Right |
| **EHS 2** | 6 | M | Hippocampal sclerosis | Right |
| **EHS 3** | 14 | F | Hippocampal sclerosis | Right |
| **EHS 4** | 17 | M | Hippocampal sclerosis/ FCD | Right |
| **EHS 5** | 26 | F | Hippocampal sclerosis | Right |
| **EHS 6** | 36 | M | Hippocampal sclerosis / DNT | Right |
| **EHS 7** | 40 | M | Hippocampal sclerosis/ FCD | Right |
| **EHS 8** | 43 | F | Hippocampal sclerosis | Right |
| **EHS 9** | 44 | F | Hippocampal sclerosis | Right |
| **EHS 10** | 50 | F | Hippocampal sclerosis | Right |
| **EHS 11** | 59 | F | Hippocampal sclerosis | Right |

Abbreviations: EHS – Epileptic Human Sample; M – Male; F – Female; FCD – focal cortical dysplasia; DNT – Dysembryoplastic neuroepithelial tumors.

**Supplementary Table 4.** Clinical information of patients who underwent autopsies.

| **SAMPLE** | **AGE** | **SEX** | **PRIMARY CAUSE OF DEATH** | **POST-MORTEM**<br>**DELAY (h)** |
| --- | --- | --- | --- | --- |
| **CHS 1** | 74 | M | Cardiorespiratory failure | 11 |
| **CHS 2** | 42 | M | Myocarditis | 10 |
| **CHS 3** | 46 | M | Bronchopneumonia | 9 |
| **CHS 4** | 53 | F | Bronchopneumonia | 7 |
| **CHS 5** | 57 | F | Aorta dissection | 8 |
| **CHS 6** | 71 | M | Bronchopneumonia | 10 |
| **CHS 7** | 72 | M | Cardiorespiratory failure | 9 |

Abbreviations: CHS – Control Human Sample; M – Male; F – Female.

## Supplementary Figures

**Supplementary Figure 1.**
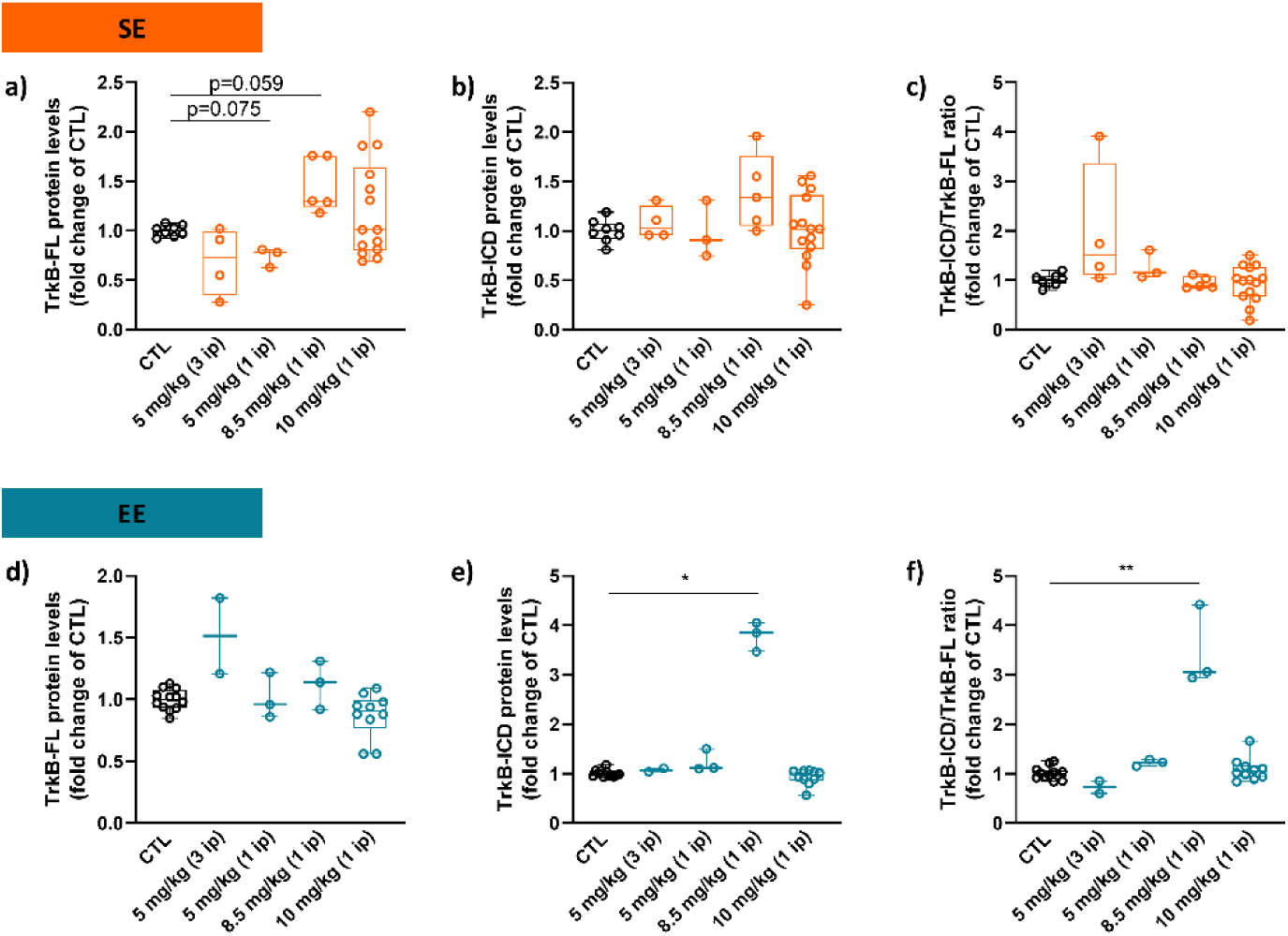
Cortical protein levels of animals treated with different doses of kainic acid (KA) and sacrificed during status epilepticus (SE) (a-c, orange) or during established epilepsy (EE) (d-f, blue) are changed depending on the KA administered dose. (a) TrkB-FL levels, (b) TrkB-ICD levels, and (c) TrkB-ICD/TrkB-FL ratio of animals sacrificed on the SE day (CTL: n=8; 5 mg/kg (3 i.p.): n=4; 5 mg/kg (1 i.p.): n=3; 8.5 mg/kg (1 i.p.): n=5; 10 mg/kg (1 i.p.): n=14). (d) TrkB-FL levels, (e) TrkB-ICD levels, and (f) TrkB-ICD/TrkB-FL ratio of animals sacrificed during EE (CTL: n=12; 5 mg/kg (3 i.p.): n=2; 5 mg/kg (1 i.p.): n=3; 8.5 mg/kg (1 i.p.): n=3; 10 mg/kg (1 i.p.): n=10). Control animals in all experiments were treated with saline (black). Statistical analysis was performed using Brown-Forsythe ANOVA and Welch ANOVA Tests (a-d) or Kruskal-Wallis test (e and f), all followed by Dunnett’s multiple comparisons tests. Results are shown as median, minimum, 25% and 75% percentile, and maximum. *p<0.05, **p<0.01, versus CTL group.

**Supplementary Figure 2.**
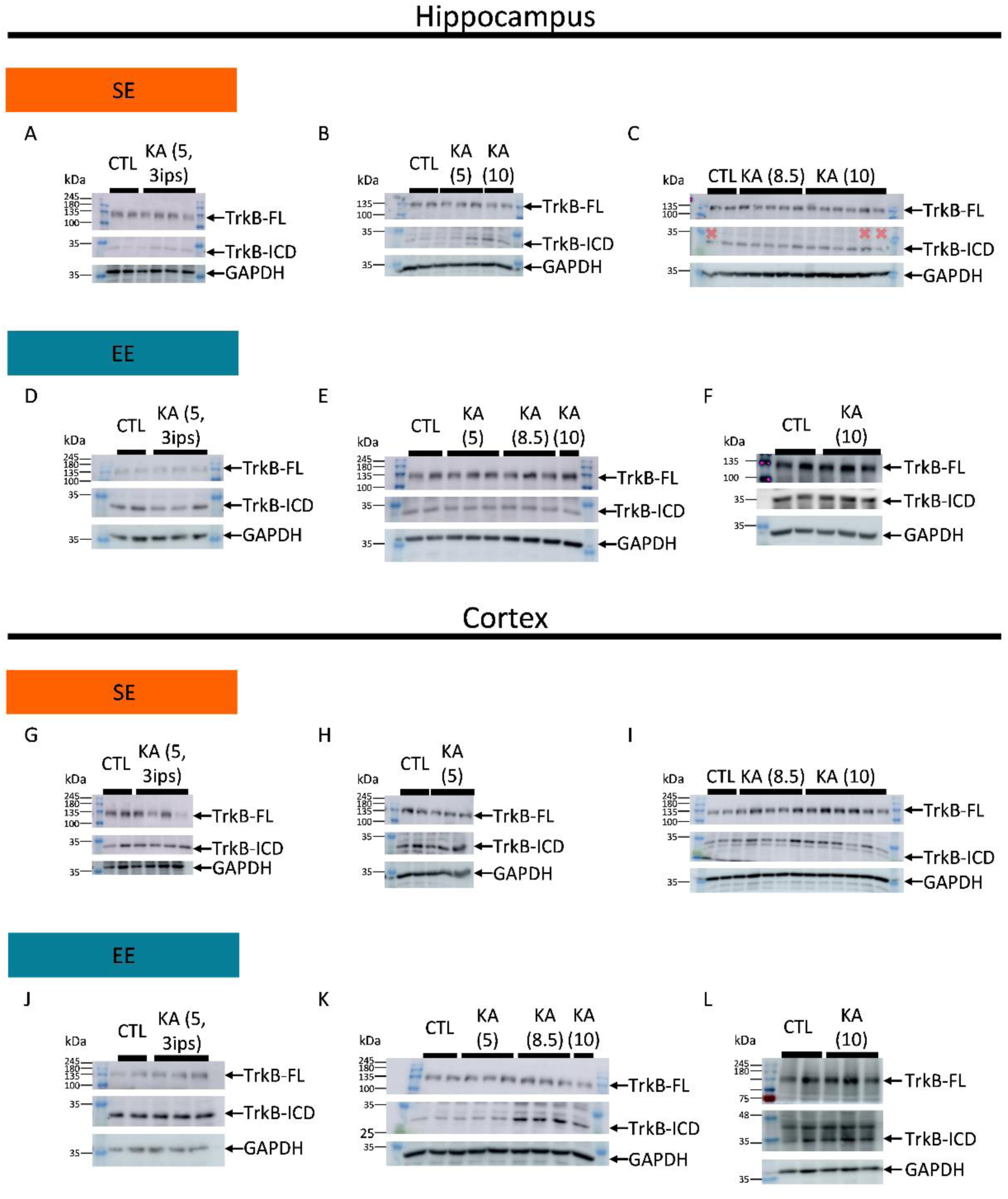
Representative western blot bands of TrkB-FL (∼145 kDa), TrB-ICD (∼32 kDa), and GAPDH (∼37 kDa) from samples of the hippocampus and cortex of animals injected with different dose regimens of kainic acid (KA) and sacrificed during status epilepticus (SE) or during established epilepsy (EE). X denotes samples excluded from the analysis.

**Supplementary Figure 3.**
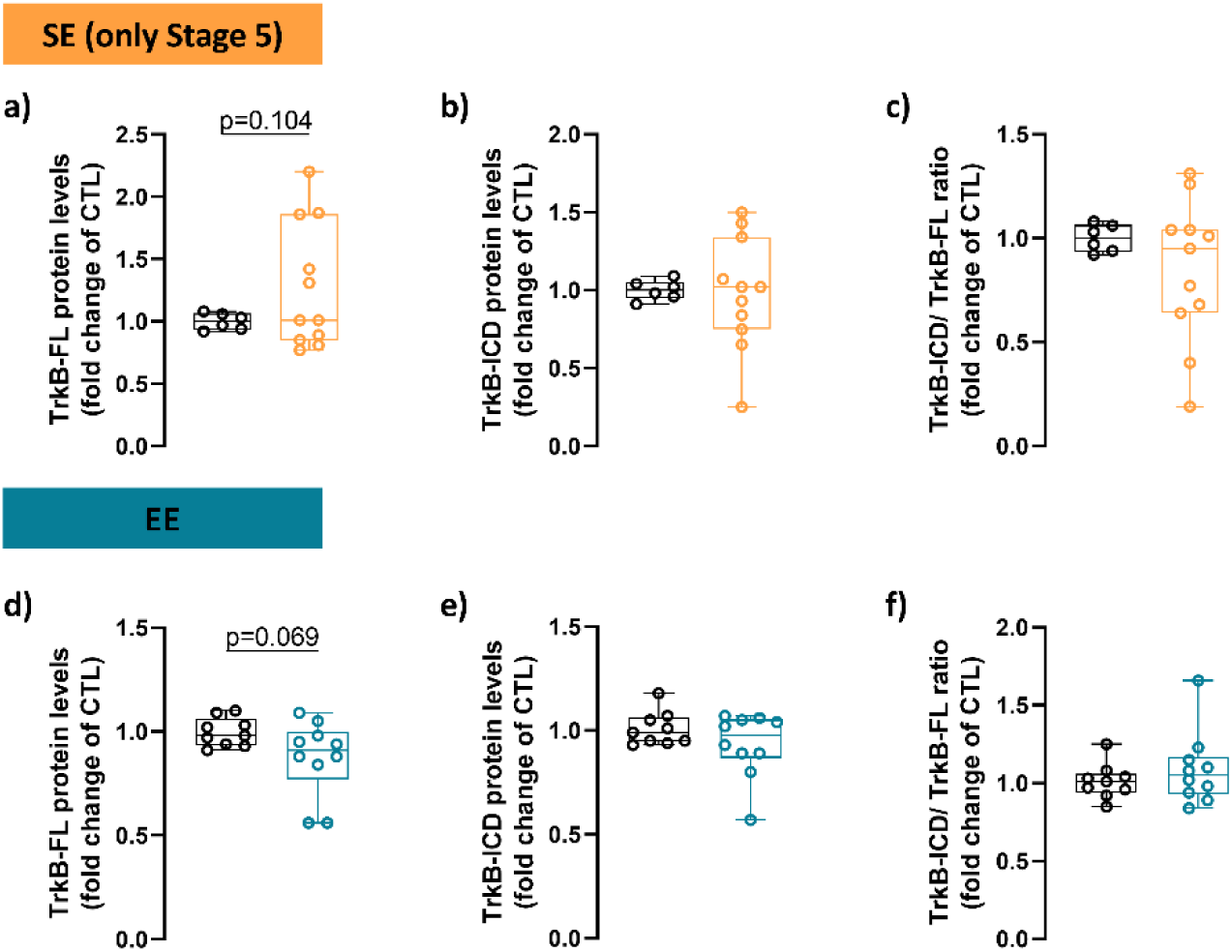
Cleavage of TrkB-FL does not occur in the cortex of animals treated with 10 mg/kg of kainic acid (KA) and stage 5 seizures during status epilepticus (SE) (a-c) or during established epilepsy (EE) (d-f). (a) TrkB-FL levels, (b) TrkB-ICD levels, and (c) TrkB-ICD/TrkB-FL ratio from the cortex of control animals (black, n=6) and animals treated with 10 mg/kg of KA that experienced stage 5 seizures (orange, n=11) during SE. (d) TrkB-FL levels, (e) TrkB-ICD levels, and (f) TrkB-ICD/TrkB-FL ratio from the cortex of control animals (black, n=9) and animals treated with 10 mg/kg of KA (blue, n=10) during EE. Statistical analysis was performed using an unpaired t-test with Welch’s correction. Results are shown as median, minimum, 25% and 75% percentile, and maximum.

**Supplementary Figure 4.**
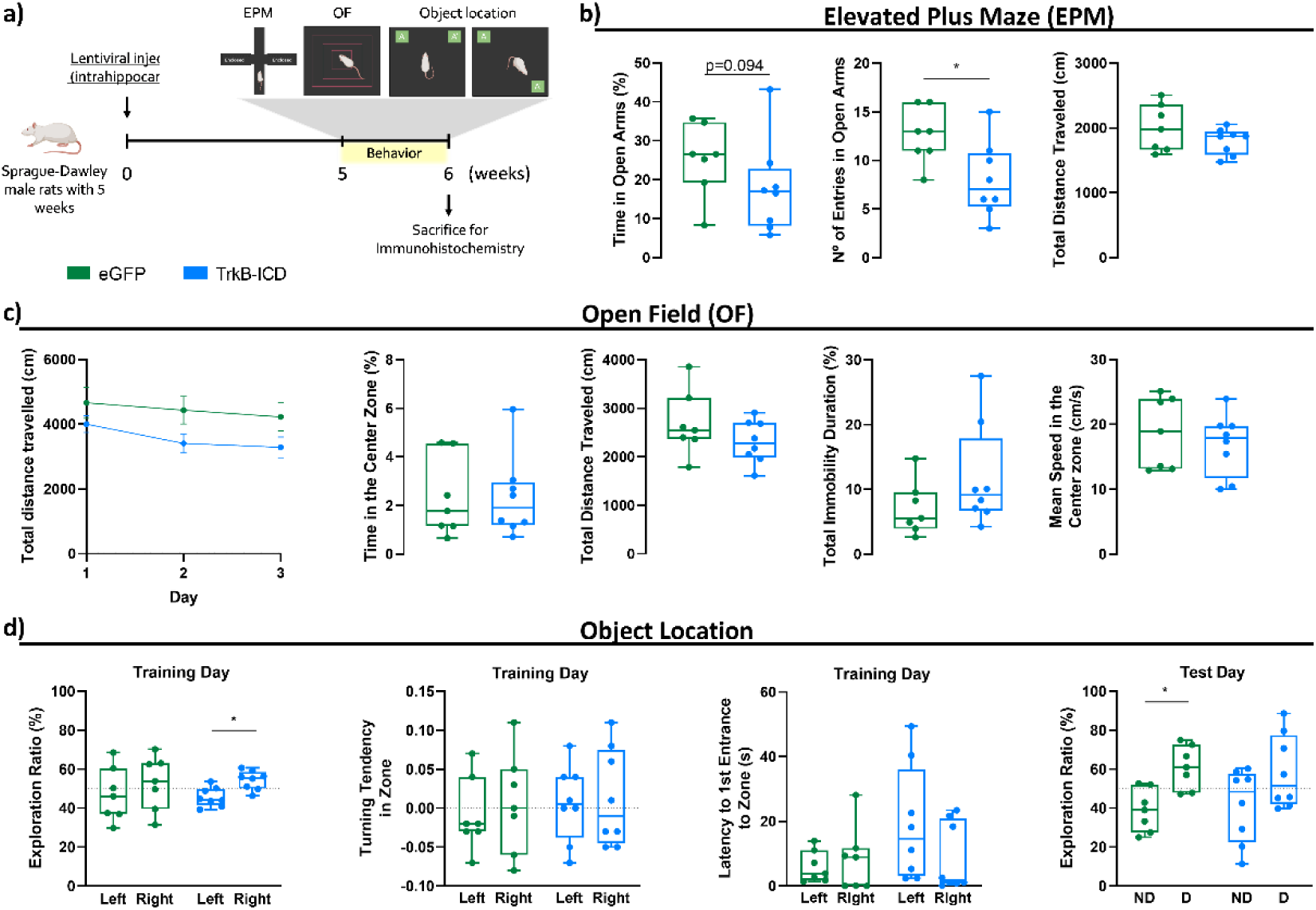
Sprague-Dawley rats overexpressing TrkB-ICD present anxiety-like behavior and long-term memory impairment. (a) Experimental design of animals injected with eGFP and TrkB-ICD lentiviral vectors. Lentiviral vectors were injected 5 weeks before the behavioral test battery. (b-d) Evaluation of (b) elevated plus maze, (c) open field test, and (d) object location test of animals overexpressing eGFP (green, n=7) or TrkB-ICD (blue, n=8). Statistical analysis was performed using an unpaired t-test with Welch’s correction (b,c), Mann-Whitney test (in the time in open arms of b, and the total immobility duration of c), or paired t-test (d). Results are shown as median, minimum, 25% and 75% percentile, and maximum. *p<0.05. ND, No-displaced. D, displaced.

**Supplementary Figure 5.**
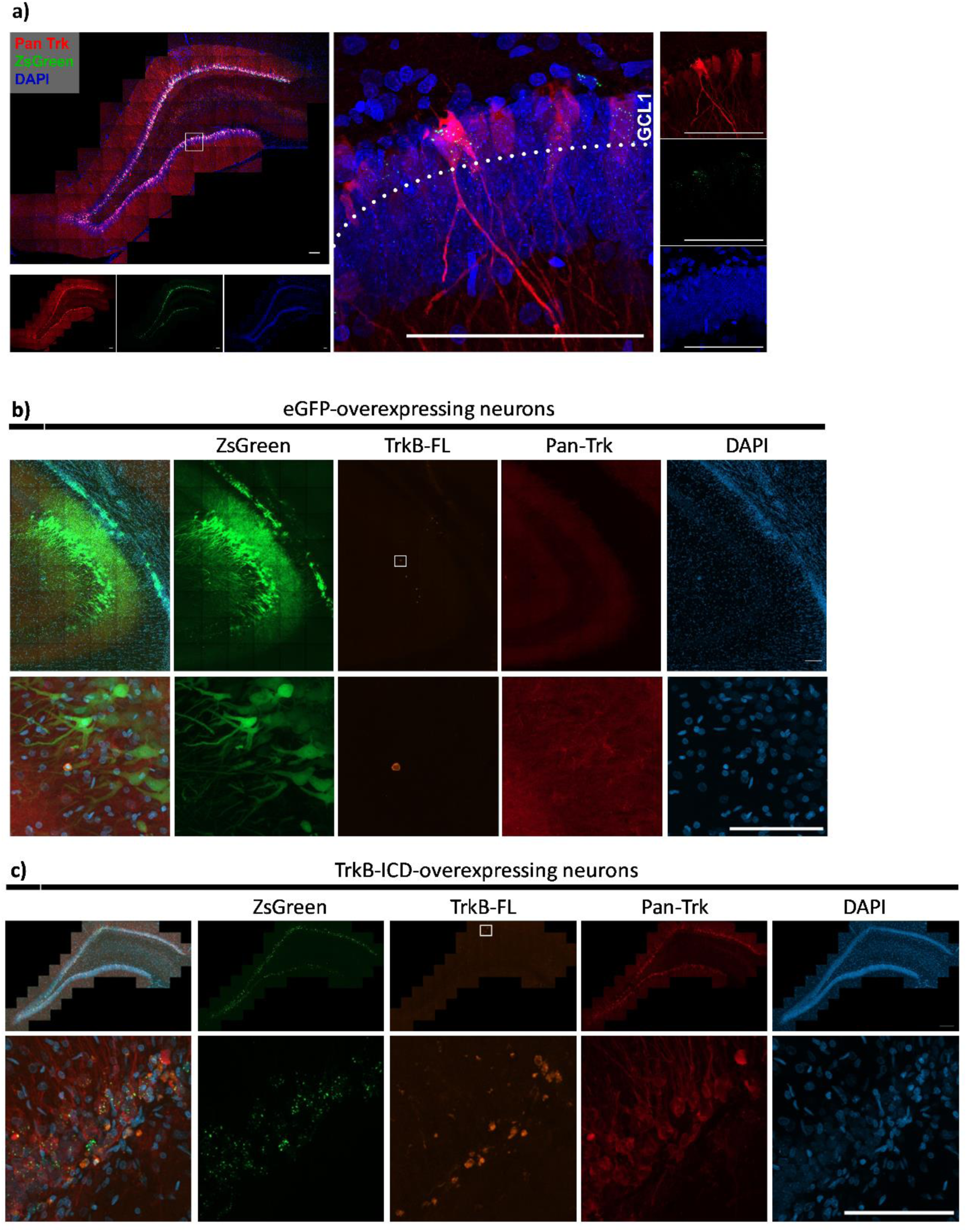
TrkB-FL and TrkB-ICD expression in the hippocampus of Sprague Dawley rats injected with eGFP and TrkB-ICD lentiviral vectors. (a) TrkB-ICD overexpression in the hippocampi of Sprague-Dawley rats. Representative immunofluorescence images show TrkB-ICD overexpression in the dorsal dentate gyrus (DG) of a rat injected with the TrkB-ICD lentiviral vector. The white square in the image indicates the area from which the right panel images were captured. Pan-Trk and ZsGreen are localized in neurons of the innermost layer of the granule cell layer 1 (GCL 1) within the DG. In all images, red staining represents TrkB-ICD (probed with anti-Trk C-terminal tail antibody, Pan-Trk), green staining indicates ZsGreen fluorescent protein (unprobed), and blue staining depicts cell nuclei (probed with DAPI staining). Images from (a) were acquired from Cell Observer SD (ZEISS) (left panel) or Confocal 980 (ZEISS) (right panel). Scale bar: 100 μm. (b) Representative immunofluorescence images of TrkB-FL and TrkB-ICD expression in the CA3 of animals injected with eGFP lentiviral vector. (c) Representative immunofluorescence images of TrkB-FL and TrkB-ICD expression in the dentate gyrus of animals injected with TrkB-ICD lentiviral vector. In images from (b-c), green staining indicates ZsGreen fluorescent protein (unprobed), orange labels TrkB-FL (probed with anti-TrkB N-terminal tail antibody), red labels TrkB-ICD and TrkB-FL (probed with anti-Trk C-terminal tail antibody, Pan-Trkblue staining depicts cell nuclei (probed with DAPI staining). The white square in the images indicates the area from which the lower panel images were captured. Images from (b-c) were acquired from Cell Observer SD (ZEISS). Scale bar: 100 μm.

